# Spatial second-order features predict glioma malignant transformation

**DOI:** 10.64898/2026.08.14.744974

**Authors:** Michal Polonsky, Jonathan J. Fox, Yunrui Lu, Sheel Shah, Noa Hadas, Jina Yun, Brian A. Williams, Barbara J. Wold, Richard G. Everson, Matt W. Thomson, Long Cai

**Author notes:** These authors contributed equally (Michal Polonsky, Jonathan J. Fox, Yunrui Lu, Sheel Shah).

## Abstract

Isocitrate dehydrogenase (IDH) mutant gliomas often transform from low to high grade aggressive tumors. The genetic and molecular drivers of this Malignant Transformation (MT) are poorly understood, and predicting whether a patient will undergo MT is an unmet challenge of high clinical relevance. To stratify patients by MT risk, we applied integrated spatial DNA and RNA profiling to biopsies from 18 retrospective glioma patients which will either remain stable, undergo MT or have already transformed. The resulting dataset consisted of >600,000 single cells, measuring 962 DNA loci, 1150 RNAs, and their spatial locations. Using this dataset, we found that genetic copy number alterations (CNAs), cell types compositions, and cellular neighborhoods did not predict MT. Instead, second order effects i.e. pairwise interactions, are highly predictive of future transformation. First, we identified abnormal chromosomal contact patterns that clearly separate future stable versus future MT samples. Second, we identified 24 ligand-receptors (LR) pairs cross-expressed in neighboring cells as the main molecular factors predictive of transformation and recurrences. We then validated a cross-expressing pair of ENPP2-LPAR1 interactions with a separate cohort of patient samples. In addition, using the LR+ cell pairs as an anchor, we identified cell signaling-specific gene expression programs that can predict from bulk or single cell RNAseq data the time to recurrence. We used a cell-interaction-based foundation model (CIFM) optimized on the spatial RNA data in forward simulations and identified potential myeloid signaling factors involved in MT. Lastly, we analyzed the effect of detection sensitivity on the ability to capture pertinent LR+ neighboring cells by down-sampling transcript and showed that the ability to detect cross-expressing signaling LR transcripts (typically <10 copies per cell) decays rapidly with lower sensitivity, but is more robust to down-sampling of the areas of the tissue imaged. The importance of second-order features suggests that increased depth and dimensionality of data on a smaller quantity of samples can provide valuable insight, and that high-sensitivity and multiple-modalities spatial approaches can help identify markers to risk-stratify patients, aid in therapeutic decision making, and uncover potential therapeutic targets.

## Introduction

Diffuse gliomas, the most common brain malignancy ^1^, are defined by isocitrate dehydrogenase (IDH) mutation status, with IDH-wt defined as glioblastoma (WHO grade 4) and IDH-mut tumors which are further sub-divided into astrocytoma and oligodendroglioma subtypes (WHO grades 2– 4)^2^. Approximately half of all patients with IDH-mut low-grade gliomas will experience recurrence within 5 years^3^ and approximately 30% of recurrent tumors will transform to a high grade tumor ^4^. Patients whose tumors undergo malignant transformation (MT) carry an extremely poor prognosis with an overall survival of < 2 years in most cases ^5^. Despite the high prevalence of MT, this process is poorly understood and predictive markers for progression and transformation are primarily demographic or radiographic such as age greater than 45, subtotal resection, tumor size greater than 3 cm, and no radiotherapy^6^. While all tumors will likely progress to MT if left untreated, identifying which patients are likely to progress rapidly after standard of care therapy would help clinicians guide further therapy, surveillance imaging intervals, and conversations regarding prognosis. The current standard of treatment involves aggressive surgical resection along with adjuvant chemotherapy and radiation, which are applied without attempting to predict tumor progression to MT ^7^.

A series of landmark studies using genomics and single-cell transcriptomics has established that gliomas are organized into a set of malignant cell states shaped jointly by genetic alterations^8–10^, intra-tumor heterogeneity and cellular plasticity^11–13^ as well as interactions with specific myeloid populations^14–16^ as drivers of glioma aggressiveness. While being foundational to our understanding of gliomas, most of these studies focused on longitudinal comparisons across tumor grades and treatment outcomes, and our understanding of the molecular processes and cellular interactions within the low grade tumors leading to progression remains limited.

To address this knowledge gap, we study glioma progression and seek predictive signatures for MT by profiling the tumor genome and the transcriptome with spatial context preserved to identify cellular interactions. We reasoned that this approach would allow us to identify first-order (cell autonomous) molecular and cellular signatures and, additionally, second-order signatures that depend on interactions amongst cells and within nuclei. We exploited the multimodal ability of our seqFISH spatial transcriptomic platform^17,18^ to probe the expression of 1150 genes at the RNA level together with 962 DNA loci in the same cell at single cell and single molecule resolution. The RNA expression data identified cell types and expression programs, while the DNA data revealed genomic patterns in a cell type specific fashion. The first-order features included cell type compositions, single cell gene expression profiles and genomic copy number alterations (CNAs), while the second-order features involving interactions of pairs of molecular features, such as ligand and receptors cross-expressed on neighboring cells, or alterations in the pairwise contact between genomics loci. This high sensitivity spatial genomics data is well-suited to capture second-order features, and we were able to identify molecular predictors to help identify tumors that are more likely to take an aggressive course and transform rapidly.

## Results

### Multi-layered molecular characterization reveals heterogeneity in gliomas

Our cohort consisted of 36 samples obtained from 18 IDH-mut patients and three IDH-wt glioma patients. The IDH-mut cohort included samples from patients that did not undergo MT based on MRI findings or pathology results of repeat resections in the follow up period (future-stable, n=7 patients), patients which did undergo MT based on radiographic and/or pathology findings (future-MT, n=8 patients) and transformed recurrences (transformed, n=7). Four patients had matched initial-MT and transformed recurrences, two patients had a low-grade recurrence and the rest were not matched. IDHmut patients were characterized clinically as oligodendrogliomas (Oligos) based on chromosome 1p/19 co-deletion, and astrocytomas (Astros) with no 1p/19q deletions. We subjected all samples to RNA seqFISH analysis, and subsequent multiplexed DNA-FISH was performed on 30 samples out of the cohort (Figure 1A). Single-nucleus RNAseq (snRNAseq) was performed on 10 samples out of the cohort. For spatial RNA detection, we used a custom gene panel targeting 1150 genes, comprised of glioma specific markers and pathways^19^ as well as genes involved in tumor-immune interactions and other general tumor pathways such as epithelial-to-mesenchymal transition (EMT) and angiogenesis^20,21^. For spatial DNA analysis, we used two-layered DNA-FISH^18^, modified to allow quantification of the entire genome at 3Mb resolution on large sample regions (Figure S3A–C). The resulting dataset was comprised of 695,099 cells after rigorous quality filtering, with an average of 817±682 total transcripts with 220±105 unique genes detected per cell. Matched DNA quantification was obtained on a subset of 304,079 cells (Figure 1A, Figures 2A–B).

**Figure 1.**
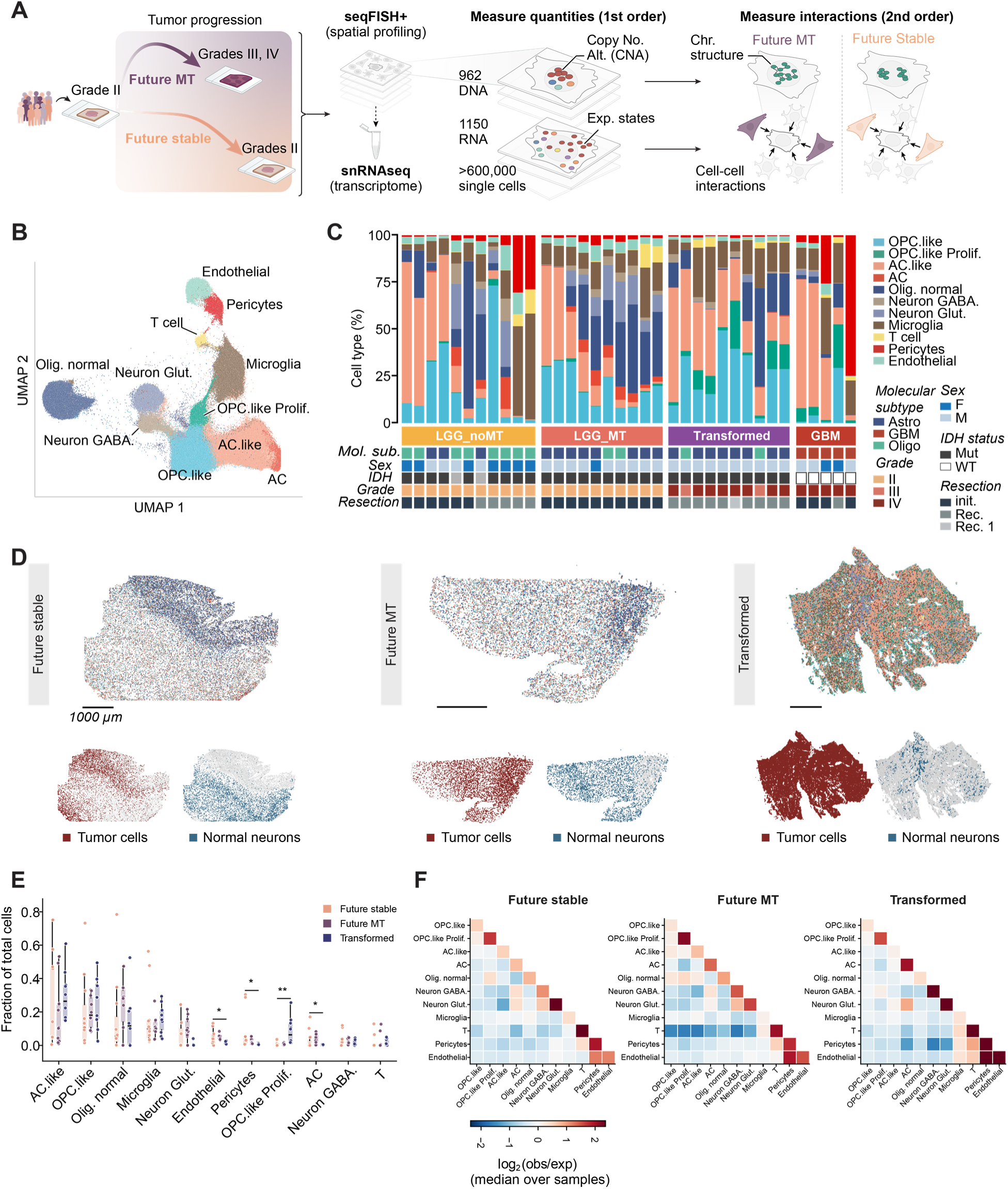
Transcriptomic and microenvironmental landscape of glioma patient biopsies. A) Experiment setup: We characterized the transcriptomic, genomic and microenvironment landscape of 36 samples from 18 IDH-mut and three IDH-wt patients using RNA seqFISH, DNA seqFISH and snRNAseq. We aimed to find molecular predictors of MT by comparing the initial resections from the IDH-mut patient cohort which either underwent malignant transformation (future MT) or did not transform during the follow-up time (future stable). B) UMAP representation of our seqFISH RNA data clustered into 11 cell types. C) Cohort composition of cell types and metadata attributes. D) Spatial representation of cell types in three representative samples from a future stable (left), future MT (middle) and transformed (right) patients. Top plot shows cell coordinates colored by the cell type as determined by spatial RNA measurements. Bottom plot shows locations of the tumor cells in red and normal neurons in blue. E) Fraction of each of the cell types within the IDH-mut samples (Kruskal-Wallis p-value; * p < 0.05, ** p<0.01). F) Proximity of each cell type to each of the other cell types within 50 µm averaged over all samples in each patient group of the IDH-mut patients. The null distribution (expected frequency) of cell interactions was calculated by random shuffling of the cell labels as described in the methods section. The heatmap values show the log_2_ of the observed frequency over the null distribution and averaged for all samples in each of the specified groups. Neither cell type compositions nor cell-type neighborhoods stratify future stable versus future MT patients.

**Figure 2:**
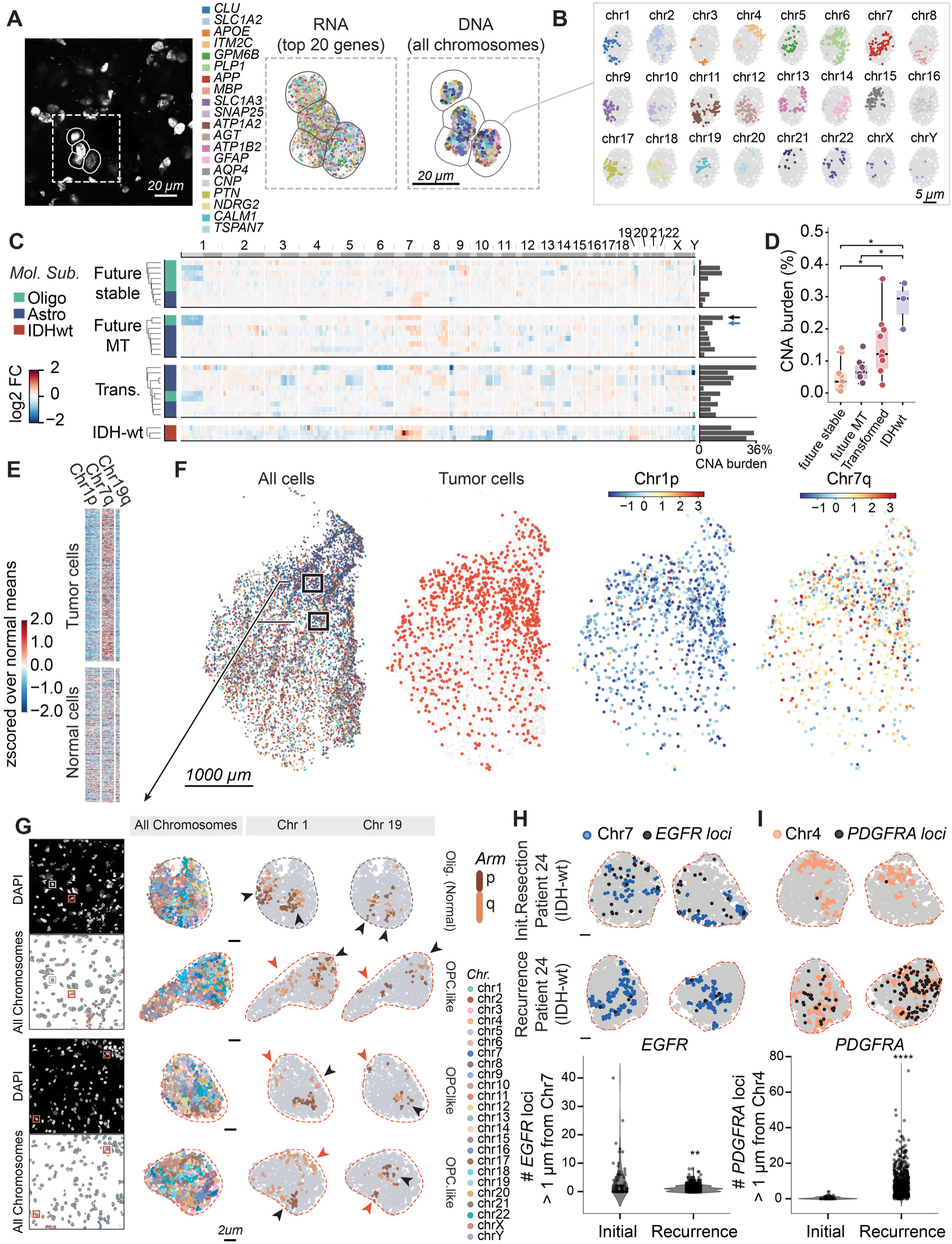
DNA-seqFISH shows common and variable CNA patterns across tumors. A) Left: zoomed in DAPI image showing a few cells from the sample presented in Figure 1D (left). Middle: spatial locations of the top 20 expressed genes detected in the labeled cells. Right: matched DNA profiles for the same cells. B) DNA locations of all the chromosomes detected from one cell. C) Mean copy number alteration (CNA) profiles of tumor cells for the IDH-wt and IDH-mut samples, measured using DNA seqFISH. CNA is defined as the log_2_ fold change of the mean DNA loci counts of the tumor cells divided by the matched normal cells within the same sample. Columns show 962 DNA loci measured in genomic order organized by chromosome number and arm (p arm, light gray; q arm, dark gray). Rows are organized by clinical outcome (IDH-wt - future-stable, future-MT, transformed; IDHmut) and hierarchically clustered within each group. Right barplots show the total CNA burden of tumor cells in each sample calculated as the percentage of loci gained or lost in tumor cells out of the total number of loci. D) Barplot comparison of the CNA burden shown in C, over the different clinical outcomes. P-values were calculated using pairwise Mann-Whitney between two groups and corrected for FDR (* p < 0.05). Black and blue arrows show patients 8 and 4 respectively and are discussed in the text. E) Heatmap of single cell CNA profiles for chromosomes 1p, 7q, and 19q in one oligodendroglioma sample comparing tumor cells and normal cells. F) Spatial maps of the same sample as in E, showing all cell type locations, locations of tumor cells, and chromosomes 1p and 7q CNA profile for each tumor cell. Boxed regions indicate zoomed in regions shown in panel G. Scale bar, 1000 µm. G) Representative images of DNA seqFISH data from the same sample showing chromosome 1p/19q codeletion in tumor cells. Left: Zoomed in regions of the sample showing DAPI staining and DNA all loci detected by DNA seqFISH colored by chromosome. Boxes indicate the representative normal cell (gray) and tumor cells (red) shown on the right. Right: Representative normal and tumor cell nuclei with genomic loci detected by DNA seqFISH. Chromosomes 1 and 19 are shown colored by arm (p arm, brown; q arm, orange). Arrowheads indicate homologous chromosomes where the arm is intact (black) or where there is an arm deleted (red). Scale bar, 2 µm. H–I) Detection of *EGFR* (H) and *PDGFRA* (I) ecDNA within one IDH-wt patient which had an initial resection and a recurrence (as labeled). Top plots show representative tumor nuclei from the initial and recurrent biopsies showing the locations of chromosome 7 and of *EGFR* loci (H) and of chromosome 4 and *PDGFRA* loci (I). Bottom plots show quantification of the number of *EGFR* or of *PDGFRA* loci detected outside of the corresponding chromosome territory (defined as > 1 µm of the nearest chromosome dot) for the initial tumor and the recurrence. P-values were calculated using two-sided Mann-Whitney (** p < 0.01, **** p < 0.0001).

By clustering on our RNA seqFISH single cell gene expression, we identified 11 unique cells types (Figure 1B, Figure S1A) comprised of normal neurons, oligodendrocytes, astrocytes, immune cells (microglia and T cells) and normal stromal cells (endothelial and pericytes), together with three populations of tumor cells which were identified based on expression of several tumor markers including *PDGFRA, DLL3* and *CD44.* To determine whether these populations corresponded to known glioma subtype signatures, we correlated pseudobulk gene expression of our seqFISH data to known IDH-wt gene programs established by Neftel et al^19^, as well as to single cell data of IDH-wt patients from the CZI database^22^ and to snRNAseq of IDHmut patients from the CARE consortium^14^ (Figures S1B–E). The two larger tumor populations in our data correlated each with two gene programs in both sets of sc/snRNAseq data and to the IDH-wt gene programs. One population correlated with the oligodendrocytic precursor cell-like (OPC-like) and neural progenitor cell-like (NPC-like) programs, which we abbreviated as “OPC-like” because the OPC marker *OLIG2* is prominent (Supp Fig S1F,G). The second corresponded to astrocyte-like (AC-like) and mesenchymal-like (MES-like) programs and was termed “AC-like” based on the expression of the astrocyte marker *SOX9* (Figures S1F–G). The third and smaller population consisted of proliferating tumor cells and was termed “OPC-like Prolif.” as it contains both OPC and proliferation markers, and is additionally correlated with G1/S gene program (Figure S1D).

### Cell type compositions, gene expression and cellular neighborhoods do not predict progression

Gliomas are known to be highly heterogeneous^8^ and, as expected, the cell type composition within our samples, as well as the spatial organization of the cells was highly variable across patients (Figure 1C). We did not detect significant differences between the cell type composition of future stable and future MT samples (Figures 1C and 1E). In line with previous studies ^11,12,23^, we found the OPC-like and AC-like cellular populations were represented both in oligodendroglioma and astrocytoma tumor subtypes. Interestingly, within the transformed recurrences, we identified an increase in the fraction of proliferating tumor cells (mean cell percentage of 0.5±1, 0.6±0.5 and 8.4±8.1 for future sable, future MT and Transformed; p = 0.0012), and a decrease in the fraction of endothelial cells, pericytes and normal astrocytes (for example, endothelial mean % are 4.9±4.2, 5.1±1.5 and 1.9±0.7 for future sable, future MT and Transformed; p < 0.05; Figure 1E). These findings are in line with the increased aggressiveness and invasiveness of the transformed tumors^24^, and suggest remodeling of the vasculature and normal stroma. At the single cell gene expression level, the OPC-like and AC-like cells were separated primarily by their cell type assignment and not by whether the cells originated from a MT or a stable sample (Figure S1A). Microglia, endothelial cells and pericytes, which could play important roles in tumor progression^25,26^, also did not separate in UMAP space (Figure S1A).

The samples were also highly heterogeneous in their spatial organization (Figure 1D and Figure S2), and while we did find that some samples showed discernable layered cellular organization of tumor and normal neurons (Figure 1D-right and Figure S2), most were highly diffused. We then sought to quantify the spatial interactions within the different patient categories and identify differences. We calculated the fraction of interacting cells of each type with each of the other cell types within 50 µm radius. To normalize the observed interaction frequency by the variable tissue compositions, we calculated the interaction frequency expected based only on the frequency of each cell type. This was calculated by permuting the cell type labels 500 times within each sample and calculating the null distribution of co-localization frequency. We then calculated the log_2_ (observed / null) for each sample, and averaged across all samples of the future stable, future MT and transformed samples (Figure 1E). This analysis showed that cells often sit close to cells of the same type, similar to observations in other tissues^27^, with the exception of microglia which are scattered over all cell-type interactions. Pericytes showed higher frequency of interactions with endothelial cells within 50 µm, which fits their biological role as supportive cells for the vasculature. The analysis did not identify significant changes in the overall spatial organization between the patient groups. For example, while spatial vicinity of normal oligodendrocytes and OPC.like tumor cells increased slightly in future MT samples compared to stable (mean log2FC of 0.22±0.31 and −0.29±0.44 respectively) the p-value was not significant after multiple hypothesis testing (p = 0.15). There was a modest, but not statistically significant, increase in the spatial vicinity of T cells, microglia, pericytes and endothelial cells within the Transformed tumors. The spatial vicinity increased by 1.3 between T and microglia comparing Transformed samples to future stable (p= 0.5). Similarly, T - endothelial, and microglia - endothelial spatial vicinity increased by 1.8 and 2.44 respectively (p=0.7, 0.15). This trend could reflect recruitment of the immune system following transformation, and influx of lymphocytes through the vasculature. However, these changes were mild and did not pass the significance threshold. We conclude, as measured, that neither the cell type composition nor the spatial organization of the low-grade tumor are predictive of future malignant transformation.

### DNA-seqFISH based CNAs identify known and patient specific genomic alterations

We next turned to our matching DNA seqFISH data to identify genomic changes between our patient categories. The genomic landscape of glioma has been well characterized^28,29^ and while some alterations are shared within molecular subtypes, such as 1p/19q co-deletion in oligodendrogliomas and ATRX nuclear expression loss and *TP53* mutations in astrocytomas^2^, the genetic makeup of individual tumors is highly heterogeneous^8^. Similar heterogeneity has previously been observed and attributed to selective pressures on individual clones within the tumor following initiation and during treatment responses^11,30^. Therefore, we reasoned that characterizing the genomic alterations in individual primary samples is of potential clinical significance and could infer patient outcomes.

We first calculated the genome-wide DNA CNA profile of each tumor sample in our data. The CNA profiles were defined as the log_2_ fold change of the mean DNA locus counts of the tumor cells divided by the mean of a null distribution of matched normal cells within the same sample. To derive null distributions, we measured the mean DNA loci count of normal oligodendrocytes and microglia, sampled randomly at equal numbers to match the tumor cells. This was repeated n=1000 times and the mean of the resulting distribution was used as the reference point to derive the tumor fold change, as well as an empirically obtained p-value for the significance of each CNA. CNAs detected by DNA seqFISH correlated with inferred CNAs from snRNAseq (Figure S4). In addition, we calculated the total CNA burden within each sample, defined as the fraction of CNAs with a |log_2_FC| ≥ 0.25 and a p-value < 0.001 out of the total number of detected loci.

The resulting CNA profiles identified known genomic alterations distinguishing between the IDH-mut glioma molecular subtypes (Figure 2C). As expected, almost all IDH-mut Oligo patients showed chromosome 1p/19q co-deletion^2^, while chromosome 7 amplification accompanied by chromosome 10 deletion were observed in IDH-wt patients^31,32^. IDH-mut-astrocytomas are characterized by mutations in *TP53* and *ATRX* which do not reflect in CNA. However, we did detect chromosome 9p deletion in a subset of patients, indicating *CDKN2A/B* deletion which is known to occur in these tumors and signifies malignant transformation (Figure 2C and Figure S3D). In addition to the similarities within tumor subtypes mentioned above, each patient had a unique CNA profile. For example, the oligodendroglioma sample from patient 8 (Figure 2C, Future MT panel, black arrow) had the characteristic 1p/19q codeletion but also unique deletions in 4p and 11p and an amplification of chromosome 7, whereas a second oligodendroglioma sample (patient 4, blue arrow) showed the same 1p/19q co-deletion, accompanied by 6p deletion and 8q amplification. The overall CNA burden was also highly variable between tumors, though we detected a significant increase in the CNA burden within transformed and IDH-wt samples (mean CNV burden of 5.1±4.8% for stable, 7±3.5% for MT, 14.5±9% for Transformed and 27.8±5.9% for the IDHwt samples). This is in line with previous studies which reported an increase in genomic alterations over glioma progression and following treatment ^8,9,33^ (Figure 2D). Thus, our spatial DNA analysis is able to capture known genomic alterations in glioma, reveal the heterogeneity in the genomic makeup of different tumors, and capture an increase in the CNA burden following transformation.

The CNA patterns can be directly visualized within individual cells in our samples. Figures 2E–F show the single cell CNA profiles of chromosomes 1p, 19q and 7q of the normal versus tumor cells from patient 8 (black arrow in Figure 2C). Tumor and normal cells are defined based on the single cell expression profiles. The single cell data reflects the patient-averaged CNA trace, and tumor cells show clear 1p/19q co-deletion and 7q amplification compared with the normal cells. The specific 1p/19q arm deletions can be detected from the chromosomal organization of single cells. Figure 2G shows several representative single tumor cells (outlined in red) and one normal cell (outlined in gray) from the same sample. The normal cell contains two intact homologous chromosomes of chromosomes 1 and 19 as indicated by the arm colors. Black arrows indicate chromosomes where both arms are present. Within the tumor cells however, one homologous chromosome is missing the p arm for chromosome 1 and the q for chromosome 19, indicated by a red arrow. Similarly, our spatial DNA analysis was also able to distinguish between hemizygous and homozygous deletions of *CDKN2A* which are known to occur in IDH-mut tumors (Figure S3D) ^34^. Therefore, the subcellular resolution of the spatial DNA data is able to capture single cell hemizygous deletions of chromosome arms, as well as distinguish the zygosity of specific gene deletions.

Furthermore, we can identify the presence of extra chromosomal DNA (ecDNA) of specific gene loci. ecDNA are small circular DNA fragments that act as oncogene carriers and drive tumor progression and treatment resistance in multiple tumor types including glioma^35^. We measured the physical distance between loci of two glioma associated oncogenes—*EGFR* and *PDGFRA*^36^—to their chromosome territory (7 and 4, respectively) within one IDH-wt patient from our cohort for which we had both the initial resection and a matched recurrence. *EGFR* loci tended to be present outside the chromosome 7 territory for the initial tumor but not in the recurrent tumors from the same patient (Figure 2H). This pattern was reversed for *PDGFRA*, where the initial tumor did not show extra-chromosomal localization, while the recurrence showed a dramatic increase in extra-chromosomal *PDGFRA* loci (Figure 2G). This suggests that our analysis is able to capture ecDNA that harbors glioma-related oncogenes, and provides potential evidence to tumor evolution and plasticity during progression, as the tumor cells shift from *EGFR* to *PDGFRA* ecDNA amplification which could confer treatment resistance.

### Alterations of chromosome contacts is a predictor for malignant transformation

Using principal component analysis (PCA), we found that while CNA features matched perfectly with the molecular subtype of the tumor (astrocytoma vs oligodendroglioma) but could not separate future MT from future stable samples (Figure 3A). Additional analysis of the CNA profiles beyond PCA also cannot differentiate between the future MT and future stable patients.

**Figure 3:**
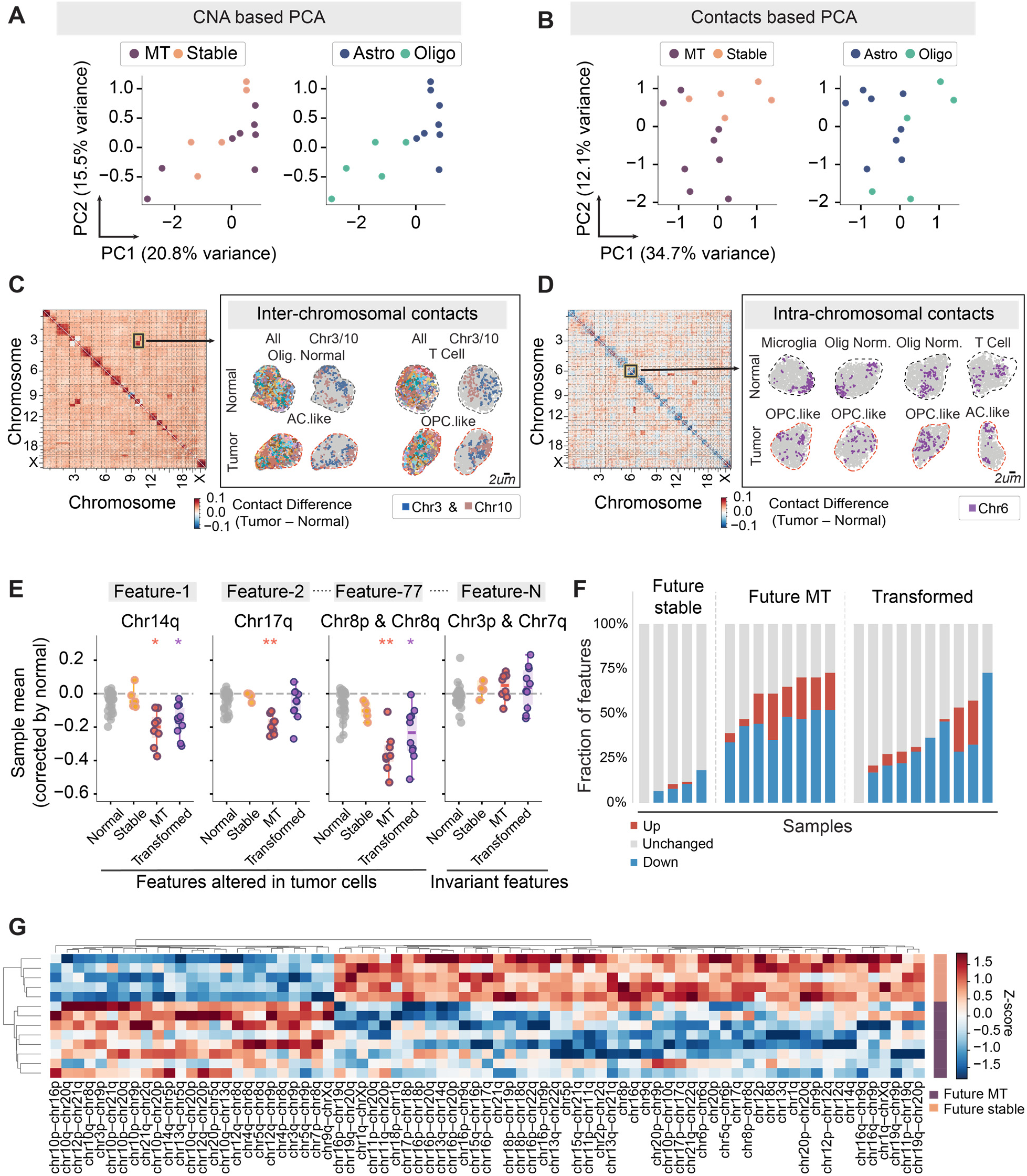
DNA second-order contact alterations predict future tumor progression. A) DNA loci-based PCA representation of the IDH-mut samples colored by state classification (future-MT and future-stable, left) and by the tumor molecular subtype classification (oligodendroglioma and astrocytomas, right). The PCA was constructed by calculating the mean CNA of each DNA loci within tumor cells in each sample, the samples were standardized and PCA was performed. B) DNA contact-based PCA on the IDH-mut samples colored as in A (left - state classification, right - molecular subtype classification). The PCA was performed as in A, except here the mean contact features were calculated for the tumor cells and binned at the chromosomal arm level in each sample. C) Left: Heatmap showing the normalized contact probability between all DNA loci in one future-stable sample. The contacts were calculated as the change in contact probability within 1um between tumor and normal cells, corrected as detained in the methods section. Inset shows enhanced contact probability between chromosomes 3 and 10 in this sample. Right: Example of normal and tumor cells within the sample showing all chromosomes, and chromosomes 3 and 10. The two chromosomes are delineated within normal cells and mixed within tumor cells. D) Left: Contact probability heatmap of one future-MT sample. Probabilities of contacts were calculated as in C. Inset shows reduction in contacts in chromosome 6. Right: Examples of normal and tumor cells from the same samples, showing chromosome 6 is more compact in normal cells compared to the tumor cells. E) Examples of statistically significant and insignificant DNA contact features at the chromosome-arm level. The plot shows the distribution of selected DNA contact features over the sample state classifications (future-stable, future-MT and Transformed). Features which were significantly different than the normal means in each category are indicated with an asterisk. Each dot represents the mean contact feature of all tumor cells within a sample, normalized by the mean of the normal cells within the same sample. We identified 77 DNA contact features which were significantly different between the sample means vs null and significantly different between the future-stable and future-MT samples. F) The number of contact features within each sample which are either up (red) down (blue) or unchanged (gray) which are 0.1 SD away from the null distribution. These were calculated for the 77 significant features found in E. G) Heatmap showing the relative abundance of each of the 77 significant features within the future-stable and future-MT samples.

We next asked whether higher-order nuclear organization of the tumor cells could differentiate between the patient categories. Recent studies have identified heterogeneity of chromosome structures within IDH-wt tumors ^37,38^. We calculated the contact frequency as the probability of proximity of each chromosome loci with another locus within 1 µm. We observed changes in both interchromosomal and intrachromosomal contacts in tumor cells compared to non-tumor cells in the same sample. In a future stable patient, we observed that chromosome 3p and 3q arms lost contact with each other and gained contacts with chromosome 10q and 10p arms, respectively (Figure 3C). These can be observed in both the contact maps (left) and in the DNA seqFISH images of individual cells (right) within each sample type. Interestingly, apparent intrachromosomal decondensation in tumor cells was observed in multiple samples, as shown in one future MT patient sample for both contact maps and single cell images (Figure 3D and Figure S5).

To quantify the chromosomal contacts differences in future-stable and future-MT samples, we performed PCA and feature-based analysis. First, we performed PCA analysis on chromosome-arm contact matrices by binning all pairwise contact frequencies between loci at the chromosomal arm level. We calculated the contact frequency matrix of each chromosome/arm pair within 1 µm, for 861 pairs out of 1035 arm pairs after removing low detection loci. We then z-scored the contact features of the cells in each sample based on the mean of the normal cells in the same sample, averaged the normalized contact matrices over all tumor cells in each of the samples, and performed PCA (Figure 3B). Here, the contact frequencies separate MT from stable samples, in contrast to the PCA analysis of CNA profiles (Figure 3A). Interestingly, the contact features-based PCA did not separate oligodendrogliomas and astrocytomas which are defined based on CNA features. This suggests that the mechanisms altering 3D nuclear organization are distinct from those relating to chromosomal deletions and amplifications.

We then sought to identify specific DNA contact features that contribute to the separation of MT from stable samples. For each of the DNA chromosome arm contact features, we calculated the sample-wise differences between chromosome contact frequency in the tumor cells compared to the mean of normal cells within the sample. After correcting for multiple hypothesis testing, we identified 77 as significant DNA contact features which differentiated between MT and stable patients (Figure 3E). We determined that the number of significantly altered chromosomal contact features are higher in future-MT than future-stable patient samples (the mean percentage of altered features was 9.4±6.7 for future stable and 60.7±12 for future MT, p = 0.004; Figures 3F–G). We calculated the fraction of features which are up, down or unchanged in each sample, by first z-scoring the features vector of each sample by its own mean, and calculated the number of features where |Z| ≥ 0.1 out of the 77 significant features. The future MT tumor cells were characterized mostly by loss of inter- and intra-chromosomal contacts together with some gains of contact (Figures 3F–G). In MT, the contact pattern destabilization is high and predictive of progression relative to the stable tumors. The quantitatively dominant contact losses potentially reflect regional domains of chromatin decondensation or other chromosomal instabilities.

Taken together, our findings suggest that the physical structures of the chromosomes, rather than chromosomal deletions or amplifications, are a key early predictor for tumors which will undergo MT, differentiating them from tumors that will remain stable.

### Signaling cell pairs predicts future progression in low-grade glioma

It has been shown that interactions with the microenvironment contributes to the aggressiveness of IDH-wt tumors^39–41^. Since cell type compositions and global neighborhoods did not stratify the future-stable vs future-MT samples, we reasoned that signaling specific interactions across neighboring cells (second-order features) could be predictive of transformation. To measure these interactions directly, we quantified the expression of known ligand and receptor (LR) pairs (obtained from the NicheNet database ^42^) present in our seqFISH gene set (770 total LR genes) over all cell types in our data. We then quantified, for each LR for a given cell-type pair, the per-sample joint frequency of cross-expression, i.e. the ligand-expressing sender cell type and the receptor-expressing receiver cell type within a 50 um proximity radius. Then, to identify LR pairs which are significantly different between the future MT and future stable samples, we first compared the resulting cross-expression fraction of each pair between our sample categories and determined significance. To correct for the false discovery rate, we shuffled the future-MT and future-stable sample labels and calculated the LR pairs that are significant after multiple hypothesis testing (Figure S6).

Our LR analysis identified a total of 24 LR interacting pairs (two-sided Mann-Whitney p<0.001; FDR=0.154) that occurred more frequently in the future MT samples compared to stable. For example, neighbor cross-expression of *ENPP2*-expressing normal oligodendrocytes with *LPAR1*-expressing tumor cells is present in a subset of oligodrendrocyte-tumor cell pairs (Figure 4A). This cross-expression is enriched in all future MT samples compared to stable ones (Figures 4B–C, 29.54 ±7% vs 5.14±4.27% respectively, p < 0.001), pointing to an oligodendrocyte-derived lysophosphatidic acid (LPA) signaling axis acting onto tumor cells in the progressing microenvironment. Increased LPA pathway activation has been observed in IDHwt gliomas ^43^ and in other tumors ^44^ and found to be central to tumor pathogenesis and progression by affecting cell proliferation, survival and differentiation and promoting EMT, cell migration, immune interactions and angiogenesis^45,46^. Notably, the total fractions of tumor cells and oligodendrocytes are heterogeneous and not statistically significantly enriched in future-MT versus future-stable samples (Figure 1C). Therefore, the interacting pair identifies a specific subpopulation of these cell types indicative of early transformation processes.

**Figure 4:**
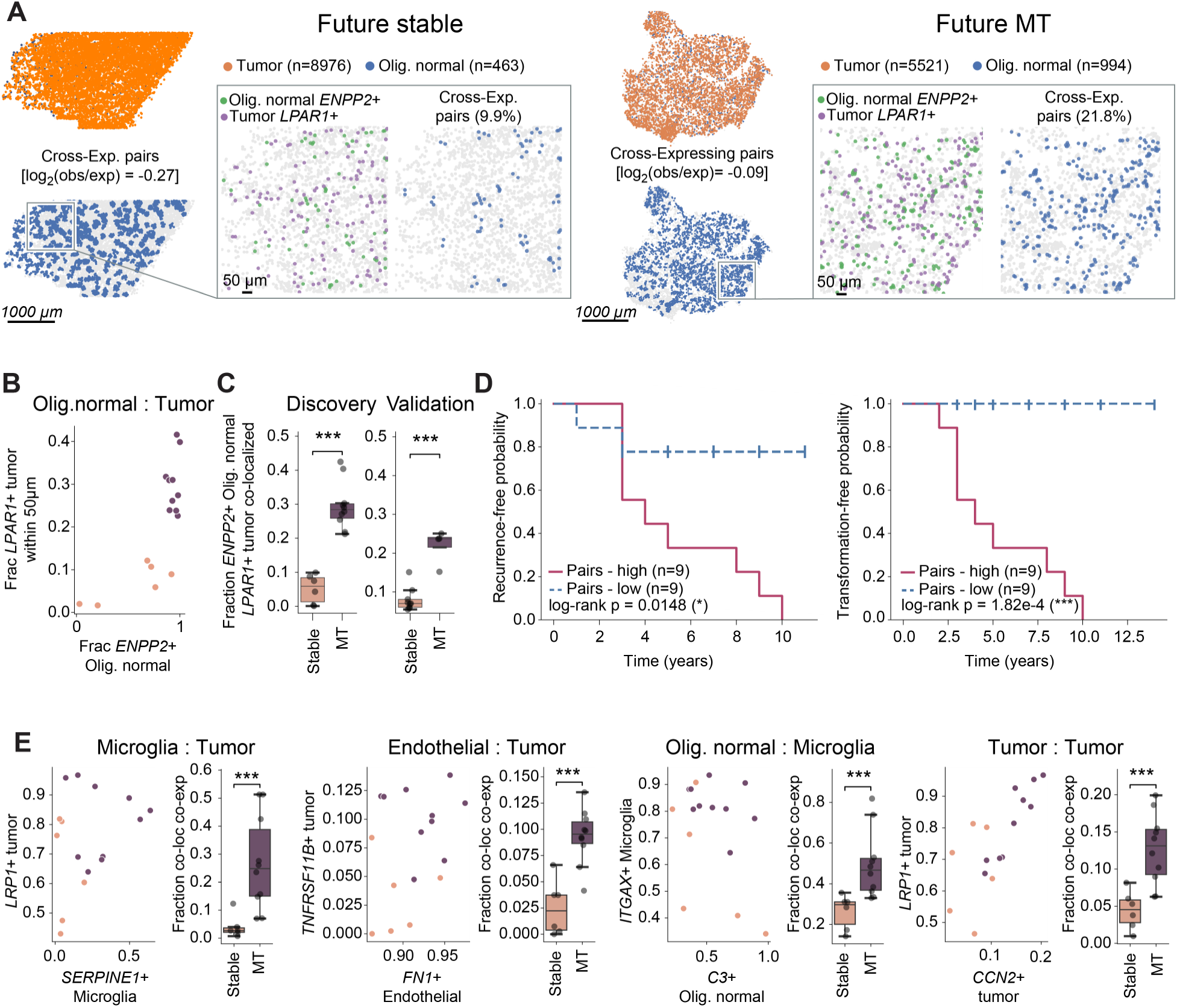
Cross-expression of ligand-receptor on cell pairs stratify MT and stable patients. A) Representative images of one stable (left) and one MT (right) tissues, showing the locations of normal oligodendrocytes and tumor cells (top, Oligo. normal and tumor cells are indicated in color) and the locations of the Olig.normal - tumor pairs found within 50um of each other (bottom). The total number of cells of each type is indicated in the figure, and the spatial enrichment of the pairs is indicated as the log2(obs/expected) was calculated as described in Figure 1F. Within each plot, a zoomed in inset shows the locations of the *ENPP2*+ Oligo. Normal and the *LPAR1*+ tumor cells highlighted in color, and the location of the co-localized and co-expressing cell pairs. The fraction of co-localized and co-expressing pairs in each sample is indicated (future stable: 9.9%, future MT: 21.8%). B) Spatial cross-expression analysis of *ENPP2*+ Oligo. Normal with *LPAR1*+ tumor cells in future-MT (n=10 samples from m=8 patients) and future-stable (n=6 samples from m=5 patients). For each pair (sender cell type to receiver cell type), the scatter plot (left) shows two independent in-pair expression marginals computed at the cell level: the x-axis is the per-sample fraction of sender cells expressing the ligand (≥1 raw transcript count) among sender cells with at least one receiver-cell type cell within 50 µm, and the y-axis is the analogous fraction of receiver cells expressing the receptor among receiver cells with at least one sender cell type cell within 50 µm. The boxplot (right) shows the per-sample cell-pair co-expression fraction, computed at the cell-pair level: the fraction of all within-50-µm (LR+ sender and receiver) cell pairs in the sample. The denominator enumerates cell pairs, a sender cell with k receiver-cell type neighbors within 50 µm contributes k pairs to the denominator. Colors indicate future MT and future stable samples. C) fractions of the *ENPP2*+-*LPAR1*+ pair. Left - fractions detected with our discovery cohort of patients. Right - same pair detected in an independent validation cohort which included 4 additional patients (1 MT and 3 stable) with a total of 13 samples from all patients. D) Kaplan-Meier curves of Recurrence free survival (left) and Transformation free survival (right) for a combined set of discovery and validation patients (n=18). The patients were stratified by the median fraction of the *ENPP2*+*LPAR1*+ : Olig. normal-tumor pairs. Patient numbers and log rank p-values are indicated. E) Spatial cross-expression analysis of four representative ligand–receptor (LR) for cell pairs, similar to B.

We then validated this finding with a separate cohort of four new IDHmut patients, three matching our future-stable definition and one future-MT. We performed seqFISH analysis on 13 sections from these patients with a reduced pool of 130 genes including cell type markers as well as *ENPP2* and *LPAR1*. The resulting single cell profiles were referenced mapped to the large seqFISH cohort to identify cell types based on our existing definitions, and the fraction of *ENPP2*-*LPAR1* pairs were quantified for each sample. Both our discovery (Figure 4C, left) and validation (Figure 4C, right) cohorts show this LR pair is significantly enriched in future MT samples. Finally, we stratified all low-grade patients from the discovery and validation cohort (n=18, future stable and future MT) based on the abundance of the pair, and generated Kaplan-Meier curves for recurrence-free probability (Figure 4D, left) and transformation-free probability (Figure 4D, right). We found that in both cases, the patients with high fractions of the *ENPP2*-*LPAR1* pair (above the median) progressed and transformed significantly faster compared to patients with low pair abundance (log-rank p = 0.0148 and 0.000182 respectively). The Cox hazard ratio was also significant for both cases (0.0127 and 0.0183) after considering the molecular subtype (Astro, Oligo) as a covariant. Thus, our LR pair analysis identified the LPA signaling axis between oligodendrocytes and tumor cells as predictive of transformation and recurrence.

### Cell signaling-specific gene programs predict recurrence time in bulk RNAseq data

Additional LR pairs relating to tumor and immune associated signaling pathways (Figure 4E, Figure S7) were identified. The complement pathway, involving *C3* and its receptor components *ITGAX* and *ITGAM* are present in several cell pairs, indicating that scavenging of lipids and cell debris is upregulated in malignant transformation. *FN1* and *CCN2* are signatures of ECM in the tissues^47^. *SERPINE1* inhibits ECM degradation and reinforces ECM^48^. *LRP1* and *LRP2* are involved in clearing of LDL and lipid droplets^49^. These LR pairs capture interactions between the microenvironment and the tumor cells implicated in EMT and microenvironmental remodeling fueling the growth and progression of various tumors including gliomas^50,51^.

Microglia participate in over half (54%) of all the significant LR pairs we detected. Myeloid subpopulations play important roles in glioma progression^26,52^ with myeloid-tumor interactions participating in IDH-wt glioma invasiveness^53^ and support tumorigenesis ^15^ and in IDH-mut gliomas. A macrophage gene program was shown to promote mesenchymal states in tumor cells of transformed tumors ^14^. We therefore asked whether we can identify cell signaling-specific gene programs which are activated in microglia. We computed the differential gene expression levels of the microglia within signaling cell pairs (‘Cross-Expressing’) compared to other Microglia that are either not cross-expressing or not in physical proximity (‘Rest’, Figure 5A). Figure 5B plots the mean delta comparing expression of microglia in a cross-expressing microglia-tumor pair with those not interacting (Rest). Interestingly, the same genes were up-regulated within cross-expressing microglia for both the MT and the stable samples, indicated by the diagonal pattern when the differential gene expression levels are compared between the two patient categories (Figure 5C). Similar patterns of cell-signaling specific gene expression are observed in other cross-expressing LR+ cell pairs (Figure S8). This finding indicates that signaling between cells is associated with a specific gene expression program, regardless of stable or MT, but increases in proportion of signaling cell pairs in future-MT compared to future-stables samples.

**Figure 5:**
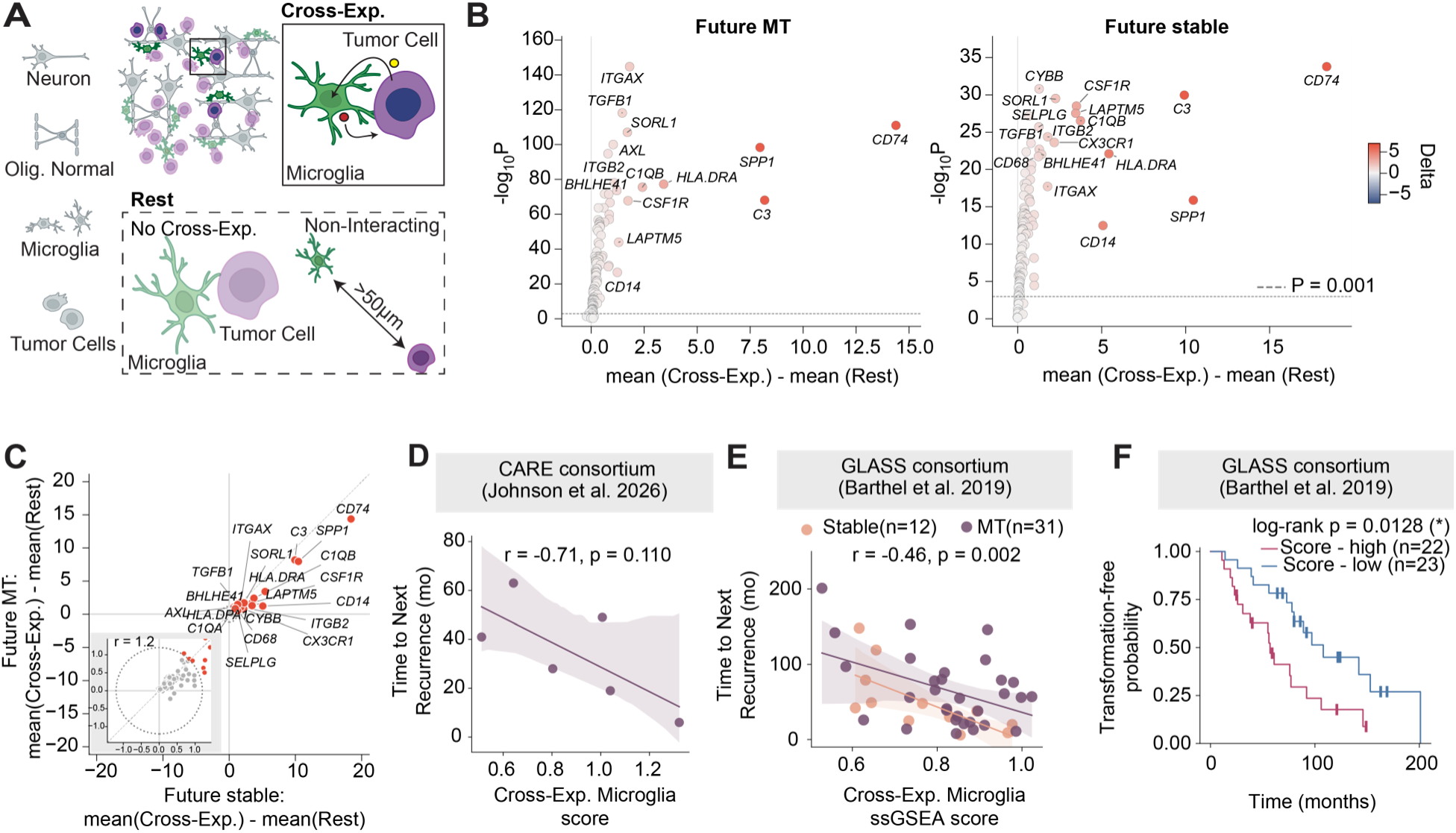
Cell signaling specific gene expression programs predict time to recurrence in existing bulk RNAseq data. A) Illustration of the analysis presented in panels B–C. Interacting cells express a specific set of genes as compared to cells either not interacting or interacting but not expressing the specific gene pairs we identified. B) Volcano plots of gene expression within microglia cells comparing microglia within LR+ pairs with tumor cells to the rest of the microglia. Left - gene expression within future MT samples, right - within future stable samples. Color indicates delta (interacting - rest). C) Same gene expression within microglia as in B, showing the delta (cross-expressing microglia – rest) for future stable on the x axis and future MT on the y axis. Zoomed in inset indicates the radius from 0,0 point that was used to identify the top co-upregulated genes. D) Correlation between the average score of microglia signaling-specific genes and time of recurrence within patients from the CARE scRNAseq dataset (Johnson et al, *Nature* 2026^14^. The score was calculated for each cell as described in the methods section, and averaged for all Myeloid cells in each sample. Only samples which were initial resections at grade II which recurred as grade III or IV are shown (Pearson r=-0.71, p = 0.11). E) Analysis of the same microglia interaction-specific genes, done on the GLASS bulk RNAseq dataset^8^. ssGSEA NES score of the gene set was calculated and correlated to time of recurrence. This dataset contained 12 samples matching our future-stable criteria (grade II recurring as grade II) and 31 matching the future MT criteria. The Pearson r and p value (−0.46 and 0.02 respectively) are calculated for all samples together. F) Transformation-free probability Kaplan-Meier curve for the same data GLASS data as in J, stratifying the patients based on the median interacting-genes score.

To determine whether these gene expression programs are informative, we analyzed the GLASS consortium bulk RNAseq dataset^8^. We found that surprisingly even in the bulk data, the selected microglia genes are anticorrelated (Pearson r = −0.46, p=0.002) with recurrence times (Figure 5E). This negative correlation was observed for patients matching our future-MT definition, as well as those matching the future-stable definition which had stable recurrences. Applying the same microglia gene expression calculation on single nuc RNAseq data set ^14^, a similar anticorrelation is observed (Pearson r = −0.71), though here only six samples matched our future-stable definition, and the p value was not significant (p = 0.11) (Figure 5D). It is interesting to note that time to second recurrence is not predicted by this signature (Figure S8), possibly because surgical resection removes the microenvironmental factors that are predictive in our analysis. Finally, we calculated the Kaplan-Meier curve of transformation-free survival times for the GLASS data. The patients’ signaling microglia score was predictive of transformation time, and initial samples with high microglia score transformed faster (log-rank p = 0.0128) (Figure 5F). In addition, signaling specific gene expression patterns in tumor cells and normal oligodendrocytes were not informative of recurrence and transformation times in the bulk RNAseq datasets, likely because the tumor genes are expressed in many cell types and are confounded in the bulk dataset, whereas the microglia genes are more distinct.

### CIFM forward simulation reveals progression dynamics latent in low-grade glioma biopsies

We next modeled the spatial microenvironments in our future-MT and future-stable samples using the Cell Interaction Foundation Model (CIFM), a graph neural network that learns how cell-cell interactions shape transcriptional states within a human tissue (Figure 6A). CIFM represents each local microenvironment of a tumor sample as a graph of transcriptional state vectors connected by spatial proximity relationships. The CIFM model acts as a generative forward tissue simulator of cell states by iteratively updating each cell’s gene expression based on its microenvironment. The 100M parameter size CIFM model was trained on 23M microenvironments across different diseased and healthy samples and fine-tuned on our complete seqFISH+ dataset using self-supervised expression reconstruction paradigm, which never sees the clinical labels.

**Figure 6:**
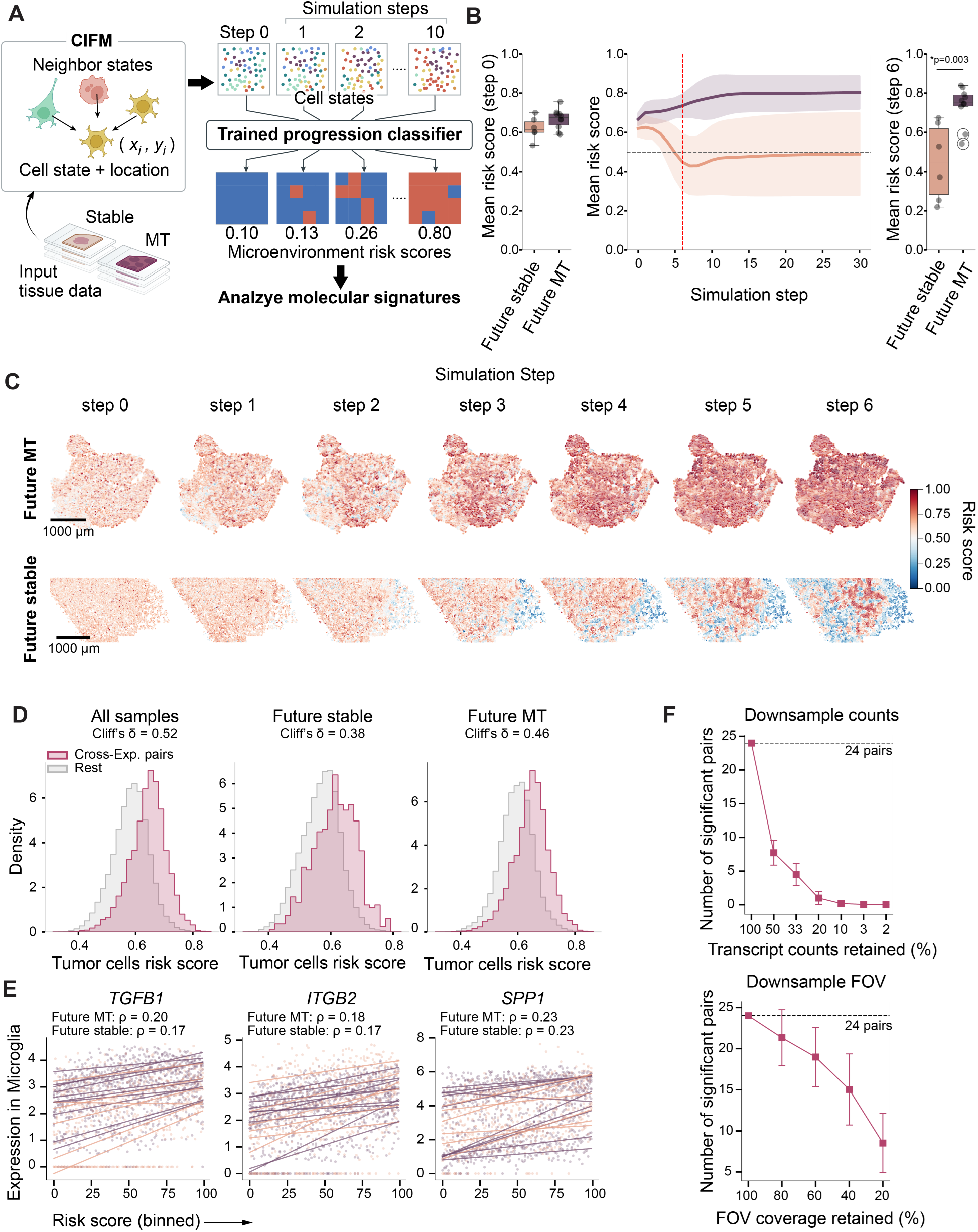
CIFM forward simulation reveals progression risk in low-grade glioma biopsies and correlates to LR+ pairs and expression of signaling specific genes in Microglia. A) Schematic of the CIFM forward simulation workflow. CIFM is a graph neural network that learns how each cell’s transcriptional state is shaped by the states and spatial coordinates (x_i, y_i) of its neighbors, and acts as a generative forward simulator of spatial single cell gene expression. RNA seqFISH data from future-MT and future-stable patient biopsies serve as input. The model is iterated forward for 30 simulation steps, producing a simulated gene expression trajectory for every cell in the tissue. At each step, a trained progression classifier is applied to the simulated cell states to assign a per-cell risk score between 0 (stable-like) and 1 (MT-like). B) Left - Per-sample mean risk scores for stable and MT patients in the initial simulation step (step 0). Middle - CIFM risk score over simulation steps. Line is the mean of each sample group, shaded area shows the 95% confidence interval across samples. Right - Per sample risk scores at step 6 of the simulation, indicated in red in the middle plot. C) Spatial maps of per-cell risk scores at the indicated simulation steps representative future-MT (top) and one future-stable (bottom) biopsy. Cell states, and therefore classifier outputs, evolve under CIFM iteration. Color bar indicates risk score from 0 (blue, stable-like) to 1 (red, MT-like). D) Distributions of the risk score of tumor cells within LR+ pairs (Olig.normal : Tumor, Red) and the rest of the tumor cells (gray). Left - all samples combined, middle - future stable samples only, right - future MT samples only. Cliff’s Delta, calculating the difference between the tumor cells within the LR+ pair and the rest are indicated on each plot. E) Expression within microglia of interaction-specific genes, plotted against normalized simulation risk scores. The simulation scores were binned to 100 bins, and the mean expression of each of the indicated genes was calculated for all microglia in each risk score bin and plotted for all samples. Linear regression lines are indicated. MT and stable data are indicated in the different colors. Mean MT and stable Spearman rho values are indicated. F) Downsampling analysis of detection of the LR+ pairs. Both plots show the number of LR+ cell-type pairs reaching Mann–Whitney significance (P < 0.001, FDR=0.154) between future-MT and future-stable samples. Top - number of pairs as a function of transcript counts retained per sample. Bottom - number of pairs as a function of field-of-view (FOV) coverage retained per sample. Count threshold = 1 transcript per cell. Error bars indicate standard deviation across replicate draws.

To apply CIFM, we embed patient samples into the model and train a classifier to discriminate MT from stable microenvironments based on the embedded seqFISH+ expression vectors (1150 genes), balanced per patient and label, with 5-fold patient-stratified cross-validation (Methods). On held-out step-0 samples, per-sample mean risk score did not robustly separate the MT/stable groups (Figure 6B). We then propagated each tissue’s expression state through 30 steps of CIFM forward simulation, using fixed cell coordinates and graph topology, and scored every cell at each step with the same classifier. The per-sample mean risk score diverged sharply across the simulation: MT samples rose above the 0.5 decision boundary, while stable samples fell below it, resulting in significant separation by step 6 (Figure 6C). The same divergence was visible at the level of individual cells: in two representative biopsies, the per-cell risk score field reorganized progressively across simulation steps, with the MT sample acquiring a uniformly high-risk distribution and the stable sample resolving into a predominantly low-risk distribution. This behavior of the model suggests that it is capturing latent variables from each sample that drive future clinical outcomes.

Signaling cell pairs are strongly correlated with risk score in the CIFM. Tumor cells involved in signaling interactions are more likely to be scored with higher risk compared to cells not expressing ligands and receptors (Cliff’s δ =0.38) (Figure 6D), indicating that the model is picking up signaling interactions as key predictors of progression. Cliff’s δ is defined as the difference between the probability that a contacting tumor cell’s risk score is higher than non-contacting tumor cell with the probability that contacting tumor cell’s risk score is lower than non-contacting tumor cell. Because CIFM assigns a risk score for each cell, MT samples can contain high risk score cells as well as low-risk score cells. We can correlate the gene expression levels across cells with different risk scores within the same sample, which avoids the sample batch effects when comparing across samples. Genes upregulated in high scoring cells are consistent with the signaling cell pair analysis, such as *TGFB1*, *ITGB2* and *SPP1* in microglia etc (Figure 6E). Ligand-receptor pairs enriched in the high-risk microenvironments in the CIFM are consistent with the LR pairs that predict progression (Figure S9). Thus, the CIFM model can identify the key molecular and spatial interactions between tumor and immune cells and differentiate MT from stable samples.

### Discovery of progression-associated interactions requires high-sensitivity spatial profiling

Ligand-receptor interactions are keys to predicting glioma outcomes and identify gene expression programs that are specific to those signaling cell pairs. Both partners could be low abundance transcripts, and the cell pair statistic requires both to be detected simultaneously in two different cells, so the probability of detecting a pair falls approximately as the square of detection efficiency. To determine whether identifying these interactions truly requires the sensitivity and scale of our measurement, we subsampled the data along two independent axes (Methods): transcript counts retained per sample, mimicking lower-sensitivity spatial-transcriptomic platforms (Figure 6F, top); and fields-of-view (FOV) retained per sample, mimicking smaller imaged tissue areas (Figure 6F, bottom). For each axis we recomputed spatial cross-expression scores across all LR pairs and counted those reaching Mann–Whitney P < 0.001, consistent with FDR analysis, between future- MT and future-stable, with 100 replicate draws per condition.

Both axes monotonically reduced the number of significant pairs, with distinct dose–response behaviors. Reducing transcript counts caused the steepest decline: the 24 lost significance below ∼20% of original counts, and the total number of significant cell pairs fell from 24 at full sensitivity to below 1 LR pairs at 20% sensitivity. FOV subsampling showed less dramatic changes, retaining 40% of imaged tissue area recovered 15 out of 24 significant pairs, indicating that the signal is broadly distributed across the tissue and that moderate reductions in the imaged area do not eliminate it.

Together, these results identify the combination of high-sensitivity in RNA detection and single cell-resolution spatial data as a prerequisite for prospective MT risk stratification of low-grade glioma. Because LR transcripts are typically expressed at low levels. Sensitivity governs whether the progression-associated interaction signature exists in the data at all: lower-sensitivity spatial measurements cannot recover it. The imaged area then determines how completely the signal can be recovered once sensitivity is sufficient.

## Discussion

Our analysis shows that the high-sensitivity spatial genomics and transcriptomics data can identify early predictors of tumor progression in IDH-mut gliomas that are potentially difficult to obtain otherwise. Our analysis draws three major observations, pointing to specific pathogenic ‘pockets’ of inter-cellular signaling within the low-grade tumors, and suggesting potential mechanisms contributing to progression. First, we were able to identify second-order features as stratifiers for glioma MT and validated the finding in a second patient cohort. At the DNA level, chromosomal interactions (contact frequencies) separated future-MT and future-stable samples (Figures 3B and 3F). Our data shows that tumor cells in the patients that are primed for MT have much more decondensed chromosomes as opposed to the tight chromosomal territories seen in normal cells. At the RNA level, our analysis identified 24 signaling ligand-receptor (LR) pairs across neighbor cells that are highly predictive of future MT. In particular, *ENPP2*-*LPAR1* pairs can stratify both recurrence and transformation time independent of other clinical data. In contrast, first-order features in our data, such as tumor cell CNA, the composition of cell types and single tumor cell gene expression profiles, or even second order features such as cell-type neighborhood compositions, are not predictive of MT, even though CNA patterns are concordant with the current classification of the molecular subtypes of gliomas.

Second, the ligand-receptors interactions we identified share commonality with those involved in the lipid-related inflammatory pathways seen in other inflammatory-fibrotic diseases ^54–57^. Lipid laden macrophages have recently been shown to promote aggressiveness in IDH-wt glioma ^58^. The LR interactions in our analysis involve lipid clearing such as complement C3 pathways and LRP1/2, as well as related fibrotic pathways (*FN1*, *DCN2*, *CCN2*, *VWF* and *SERPINE1*). Tumor-adjacent and LR cross-expressing microglia show activation of *TGFB1*, *SPP1* and *ITGB2*, corresponding to inflammatory programs that are synergistic with the complement C3 and fibrotic pathways ^59–61^. Our data suggest lipid related inflammation is a potential mechanism of progression in IDH-mut as well and identifies potential targets for early intervention.

Third, using the signaling LR pairs as anchors for analysis, we identified cell-signaling specific gene expression programs in microglia that can be used to predict recurrence time in existing bulk and single cell RNAseq data^8,14^. Myeloid cells have been shown to play key roles in glioma progression and support tumor growth^26,62^. Our analysis identifies a specific subset of microglia with pathogenic potential interacting with tumor cells. Using our foundation model, we observed that cells with higher risk scores are enriched with signaling cell pairs and have higher *TGFB1* levels, which are activated downstream in the *ENPP2*-*LPAR1* pathway.

Our analysis indicates that cell-signaling specific gene expression programs instead of cell-type specific genes are potentially powerful in discovering biomarkers and underlying biology. Detection of ligands and receptors, which are typically low abundance (<10 copies per cell), require high sensitivity measurements. These genes are not strongly correlated with a large number of other genes and do not necessarily cluster out from single cell expression analysis. In contrast, genes which identify transcriptionally distinct cellular subtypes are usually part of a larger gene program involving thousands of genes and can be clustered out even with low sensitivity, ∼5% detection rate for RNAs (5 detection events out of 100 mRNA copies in a cell) in single cell RNAseq. seqFISH sensitivity was found to be ∼50% when comparing against gold-standard single molecule FISH^17^, and a recent calibration of detection sensitivity across platforms showed a similar result for seqFISH of 271 highly expressed genes ^63^. The higher sensitivity measurements are well-suited to capture the LR interactions in the data that allows further identification of cell-signaling specific gene expression programs.

A limitation of our study is medium sample size which, especially in a disease with such heterogeneous outcomes such as glioma, restricts the number of possible comparisons and could limit statistical power. Using targeted spatial transcriptomics of 1150 RNAs, we focused on signaling ligands and receptors but cannot measure all molecular pathways. A high-sensitivity analysis with a larger panel of genes^17^ will be examined in future studies. We mitigated these limitations by validating our main findings over independent patient samples. In addition, the microglia gene programs identified from our analysis showed surprisingly strong predictive power for bulk RNAseq samples of a large patient cohort ^8^. Our study suggests that to identify predictive glioma biomarkers and pathogenic molecular pathways, sample quality and detection sensitivity should be prioritized. These principles could be generalized in other pathologies and will reveal disease-specific interactions which can then be investigated further with larger patient cohorts.

In conclusion, we used high sensitivity spatial genomics to identify ligand-receptor interactions in cell pairs, which are used as anchors to discover cell-signaling specific gene expression programs that predict imminent transformation in low grade gliomas. These predictive features based on the underlying biology of MT are integrated into the predictive CIFM model. We envision that with further investigations, the discoveries in this study will become useful clinical tools for neuro-oncologists to identify patients at high risk for MT and alter therapeutic considerations in these patients along with possible drug targets to arrest the malignant transformation process.

## METHODS

### Human Glioma Tissue Collection and Storage

Glioma tissue samples were collected from patients undergoing surgical tumor resection at the Ronald Reagan UCLA Medical Center under IRB-10-0655. Tissue was flash frozen in liquid nitrogen and stored at −80°C. Tissue samples were later moved to long-term storage at −150°C.

### Coverslip Functionalization and Flow Cell Preparation

Glass coverslips (Epredia #152460) were immersed in RNase-free water (Invitrogen #10977-023) then immersed in 100% ethanol (Koptec #V1016). Coverslips were dried for 5 minutes in a 90°C oven (Bellco Glass Inc. #7930-00110) then plasma cleaned on “high” for 5 minutes (Harrick Plasma #PDC-001). Cleaned coverslips were coated for 30 minutes at room temperature with a solution of 1% (w/v) 3-aminopropyltriethoxysilane (APTS) (Thermo Scientific #80370), 9.9 mM acetic acid (Sigma-Aldrich #A6283-500ML), and 89.1% ethanol. APTS-coated coverslips were rinsed 3 times in 100% ethanol and dried for 10-30 minutes in a 90°C oven. Dried APTS-coated coverslips were then coated overnight in a humidified chamber at room temperature with 0.1mg/mL ≥300,000 MW poly-D-lysine (Sigma-Aldrich #P1024) in 100 mM borate buffer (Thermo Scientific #28341). Coated coverslips were rinsed again with RNase-free water, dried under nitrogen gas, stored at 4°C, and used within 1 week for cryosectioning.

Flow cells were prepared using glass microscope slides with drilled inlet and outlet holes. Slides were plasma cleaned on “high” for 7 minutes, and laser cut adhesive was aligned and adhered to the slide creating a channel between the holes. Flow cells were stored at room temperature before affixing to coverslips.

### Tissue Cryosectioning, Fixation, and Pre-Hybridization Processing

Fresh frozen glioma tissue was equilibrated in a cryostat (Leica #CM3050) at −20°C for 30 minutes prior to sectioning. Tissue was affixed to a cryostat specimen chuck using optimal cutting temperature (OCT) embedding medium (Fisher Scientific #4584) and sectioned at 7 μm using a microtome blade (Sakura Finetek #4689-50). Sections were adhered to APTS- and poly-D-lysine-coated glass coverslips and immediately fixed with 4% paraformaldehyde (PFA) (Thermo Scientific #28906) in 1X PBS (Invitrogen #AM9625) for 5 minutes at room temperature. After fixation, sections were sequentially rinsed with 1X PBS and 75% ethanol before drying under nitrogen gas. Fixed sections were stored at −80°C.

Prior to sample hybridization, fixed sections were incubated overnight in 75% ethanol at −20°C. After drying the tissue under nitrogen gas, samples were permeabilized using 8% SDS (Invitrogen #AM9822) in 1X PBS for 30 minutes at room temperature. Permeabilized sections were rinsed in 75% ethanol and dried under nitrogen prior to antibody immunostaining.

### Oligonucleotide Probe Design

RNA seqFISH+ probes were designed to target a panel of 1150 gene transcripts manually curated to identify the cell types present in human glioma as well as cancer-related genes and pathways (Table S1). Barcoded probes were designed by Spatial Genomics, Inc. (Pasadena, CA) for use with the GenePS spatial genomics system (Spatial Genomics Inc., Pasadena, CA).

DNA seqFISH+ probes were designed to target loci spanning the entire human genome (GRCh38) at an average spacing of 3 Mb using a two-layer DNA barcoding scheme adapted from Takei et al., 2025^18^. The 3.10 Gb human genome was divided into 68 blocks of size 45.5 Mb, and each block was further subdivided into 15 bins of size 3.04 Mb for a total of 1,020 genomic regions. For these 1,020 genomic bins of size 3 Mb, a 150 kb sub-region was targeted for probe selection; this size was selected based on previous results from Takei et al., 2025 indicating that this size is diffraction limited. For most bins, this 150 kb sub-region was targeted to a cancer- or cell cycle-related gene selected among the genes in the 3 Mb genomic bin using ChatGPT-4o via the OpenAI API (OpenAI, San Francisco, CA). If there were no relevant genes, then the 150 kb sub-region was targeted to the center of the 3 Mb genomic bin. The genomic targets were edited to include the 46 genes on the U.S. Food and Drug Administration (FDA) list of cleared or approved companion diagnostic devices (current as of December 17, 2024)^64^: *ALK*, *ATM*, *BARD1*, *BRAF*, *BRCA1*, *BRCA2*, *CD274*, *CHEK1*, *CHEK2*, *CLDN18*, *EGFR*, *ERBB2*, *ESR1*, *EZH2*, *FANCA*, *FANCL*, *FGFR2*, *FGFR3*, *FLT3*, *FOLR1*, *HLA*, *IDH1*, *IDH2*, *KIT*, *KRAS*, *LEPR*, *MAGEA4*, *MET*, *MLH1*, *MSH2*, *MSH6*, *NRAS*, *NTRK1*, *NTRK2*, *NTRK3*, *PALB2*, *PCSK1*, *PDGFRA*, *PDGFRB*, *PIK3CA*, *PMS2*, *POMC*, *RAD51*, *RET*, *ROS1*, *TP53*. OligoMiner ^65^ was used to identify oligonucleotide probes of length 35 nt within each targeted region using balanced probe mining settings. Of the 1,020 genomic regions, 58 were not targetable due to repetitive sequences or lack of unique probes, yielding a final set of 962 genomic loci at an average distance of 3 Mb, each targeted by up to 500 probes within a maximum 150 kb sub-region. In total, there were 453,279 unique probes selected spanning the entire human genome (Table S2).

DNA seqFISH+ probes were barcoded across 14 rounds of hybridization with 3 fluorescent channels per round (647 nm, 561 nm, and 488 nm) using a two-layer DNA barcoding scheme adapted from Takei et al., 2025. The 45 Mb genomic blocks were barcoded across 9 imaging rounds using an error-correcting barcode with Hamming distance of 2 consisting of 9 pseudocolors across 2 barcoding rounds plus an additional round for error correction. Barcodes were randomly assigned per genomic block. The 3 Mb genomic bins were serially encoded using 15 pseudocolors spread across 5 imaging rounds. Each pseudocolor is assigned to a 15nt barcode sequence corresponding to a fluorescently-conjugated readout oligonucleotide assigned to a specific channel and barcoding imaging round.

The final barcoded DNA seqFISH+ oligonucleotide probe sequences were stitched together using the probe sequence generated by OligoMiner and readout sequences according to the two-layer DNA barcoding scheme. Each probe contains the following primers: forward primer (5’-GCCCCATCATGTGCCTTTCC-3’) and reverse primer (5’-CTATAGTGAGTCGTATTACCGGCC-3’). The reverse primer represents the T7 RNA polymerase binding site for *in vitro* transcription, which allows for probe circularization by hybridization and ligation of a splint oligonucleotide (5’-CATGATGGGGCGCGTAGAGTG −3’) complementary to the 5’ end of the forward primer and a 3’ universal sequence (5’-CACTCTACG-3’). The final probe sequence was assembled as follows: 5’-[Forward Primer]-[Bin Readout]-CTATAC-[Block Readout A]-CAA-[Locus-Specific Probe Sequence]-AAC-[Block Readout B]-CTATAC-[Block Readout C]-AAC-[Universal Ligation Sequence]-[Reverse Primer]-3’. Barcoded DNA seqFISH oligonucleotide probe pools were synthesized by Twist Bioscience (South San Francisco, CA) with a yield of >0.2 fmol average per oligonucleotide.

DNA seqFISH probes were also designed to target 9 additional glioma-relevant genes (*CCND2*, *CDK4*, *CDK6*, *CDKN2A*, *CDKN2B*, *EGFR*, *MAF1*, *NFKB1*, and *PDGFRA*). OligoMiner was used to select 500–1000 probes per gene within a 150 kb window. The final probe sequence was assembled as follows: 5’-[Forward Primer]-[Readout Sequence]-CTATAC-[Readout Sequence]-CAA-[Locus-Specific Probe Sequence]-AAC-[Readout Sequence]-CTATAC-[Readout Sequence]-AAC-[Universal Ligation Sequence]-[Reverse Primer]-3’. Probes for these 9 additional DNA targets were synthesized by GenScript Biotech (Nanjing, China).

Additional probes were also designed against 4 repetitive genetic elements (45SrDNA, alpha satellite DNA, *Alu* elements, and *LINE-1*) and 2 additional RNA targets (ITS1 in rRNA and *MALAT1* lncRNA). Repetitive genetic element and additional RNA target probe final sequences were assembled as follows: 5’-[Locus-Specific Probe Sequence]-AC-[Readout Sequence]-3’. These probes targeting repetitive genetic elements and additional RNA targets were synthesized by Integrated DNA Technologies (Coralville, IA).

### Probe Pool Generation

Barcoded probes were first amplified by limited cycle PCR. Probes were resuspended in water at 100 ng/μL and shaken at 9 Hz in a 37°C ThermoMixer (Eppendorf #5436) for 5 minutes to fully dissolve. The PCR reaction mixture was prepared by mixing 1 uL resuspended oligonucleotide pool, 50 μL each of 10 μM forward and reverse primers (Integrated DNA Technologies), 500 μL KAPA HiFi HotStart ReadyMix (Roche #7958927001), and 399 μL water. The reaction mixture was split into 40 tubes and run on a thermocycler (Bio-Rad #1851148) with the following settings: initial denaturation at 98°C for 4 minutes; 13 cycles of denaturation at 98°C for 20 seconds, annealing at 69°C for 20 seconds, and extension at 72°C for 20 seconds; and a final extension at 72°C for 2 minutes. The PCR product was purified using the QIAquick PCR purification kit (QIAGEN #28104) using 1 column per 4 PCR reactions with a final elution in 30 μL water per column. The concentration and purity of dsDNA was confirmed by spectrophotometry using a NanoDrop One (Thermo Scientific).

The probe pool was further amplified using *in vitro* transcription to produce many copies of RNA from each strand of DNA. The number and volume of reactions was determined using 0.5–1 μg DNA/reaction and 30 μL per reaction. The final *in vitro* transcription reaction mixtures were prepared to contain 0.5–1 μg DNA/reaction, 5,000 units/mL T7 RNA polymerase (New England Biolabs #M0251S), 1X RNAPol reaction buffer (New England Biolabs #B9012S), 10mM each rNTP (Thermo Scientific #R0481), 1 U/μL RNasin Plus RNase inhibitor, and 2.5 U/mL inorganic pyrophosphatase (New England Biolabs #M0361). Tubes were incubated in a 37°C thermocycler for 16–18 hours.

Next, the RNA was immediately reverse transcribed to generate the final ssDNA probes. Reverse transcription reactions were prepared to contain 1.5 μM RNA, 37.5 μM forward primer, 0.4U/μL RNasin Plus RNase inhibitor, 1 U/mL inorganic pyrophosphatase (New England Biolabs #M0361, 1X RT Buffer (Thermo Scientific), 3 mM each dNTP (Thermo Scientific #R0182), and 1 μL/nmol RNA Maxima H Minus Reverse Transcriptase (Thermo Scientific #EP0752). Tubes were incubated in a thermocycler with the following settings: 55°C for 2 hours, 50°C for 2 hours, and 85°C for 5 minutes. Next, alkaline hydrolysis was used to remove RNA by adding 1M sodium hydroxide (Macron Fine Chemicals #7708-10) for a final concentration of 250 mM and incubated at 65°C for 20 minutes. The solution was neutralized by adding 1M acetic acid (Sigma-Aldrich #A6283-500ML) for a final concentration of 250 mM. A neutral pH was confirmed using pH test strips (MilliporeSigma #M1095330001) and adjusted accordingly with 1M sodium hydroxide or 1M acetic acid. Gel electrophoresis was performed to confirm probe size and purity. The gel consisted of 2% agarose (VWR #0710) in 1X TBE (Invitrogen #AM9863) with 1:10,000 SYBR Safe (Thermo Scientific #S33102). ssDNA probes were mixed with DNA loading dye (Thermo Scientific #R1161), loaded alongside a 50 bp DNA ladder (New England Biolabs #N0556S), and run in 1X TBE in a horizontal electrophoresis chamber (Thermo Scientific #B1A-BP) at 150V for 30–40 minutes (Bio-Rad #1645050) before observation and imaging under blue light transillumination (Invitrogen #G6600).

To purify and concentrate ssDNA probes, 0.1 volumes of 3M sodium acetate (Invitrogen #AM9740) and 4 volumes of ethanol were added to each tube and allowed to precipitate at −20°C overnight. Probes were pelleted by centrifugation at 16,000 x g for 30 minutes at 0°C. Supernatant was removed, freshly made 75% ethanol was added without disturbing the pellet, and tubes were centrifuged at 16,000 x g for 5 minutes at 0°C. Supernatant was removed, and the pellet was allowed to air dry. Probes were resuspended and pooled in 1500 μL water, and the concentration was measured using the Qubit ssDNA assay kit (Invitrogen #Q10212) with on a Qubit 4 fluorometer (Invitrogen). Finally, probes were aliquoted and lyophilized using a freeze dryer (Labconco #710202000) to produce a final working concentration of 0.75 nM per oligo when resuspended.

### Readout Probe Synthesis and Conjugation

Readout probes (15 nt) were designed by randomly generating sequences with 40–60% GC content and screened for specificity against the mouse transcriptome using BLAST. To minimize cross-hybridization, probes containing ≥10 nt consecutive matches were excluded. The reverse complements of selected readout sequences were incorporated into primary probes according to the barcode design.

For synthesis, 5’ amine-modified DNA oligonucleotides (Integrated DNA Technologies) were conjugated to Alexa Fluor 647, Cy3B, or Alexa Fluor 488 NHS esters in 0.5 M sodium bicarbonate buffer and incubated overnight (>8 h) at 37°C. Labeled probes were purified by ethanol precipitation, HPLC, and column cleanup, then quantified by Nanodrop and diluted to a 500 nM working stock. Probes were stored at −20 °C.

### RNA seqFISH+ Hybridization and Imaging

After immunostaining, the samples were hybridized for RNA seqFISH+. RNA seqFISH+ primary probe hybridization solution was prepared by Spatial Genomics, Inc. Additional probes targeting *MALAT1* and ITS1 were added to the hybridization buffer at a concentration of 4 nM/oligo. The probe hybridization solution was denatured on a 90°C heat block (Fisherbrand #88860022) for 3 minutes then cooled for 1 minute at room temperature. The sample was washed 3 times with primary wash buffer (Spatial Genomics #10200005) before adding the probe hybridization solution and incubating at 37°C for 48–72 hours in a humidified chamber. After incubation, the sample was incubated in 70% ethanol for 1 minute at room temperature then dried under nitrogen gas before adhering a flow cell. Next, the sample was washed 3 times for 30 minutes with primary wash buffer at 37°C. Finally, the sample was washed 3 times with rinse buffer (Spatial Genomics #10100001).

Prior to seqFISH+ imaging, the RNA integrity in the sample was first confirmed by a colocalization assay. This assay uses fluorescently-conjugated readout probes targeting different regions of a housekeeping gene in the RNA seqFISH+ probe set to determine whether the RNA is intact. First, validation decode probes (Spatial Genomics #40000008) were added to the sample and incubated for 15–20 minutes in the dark. The sample was then washed 3 times with 10% (v/v) formamide, 0.1% triton X-100, 4X SSC wash buffer and incubated for 5 minutes at room temperature in the dark. Next, the sample was washed with 4X SSC and incubated with 3 μg/mL DAPI (Sigma-Aldrich #D8417) in 4X SSC for 3 minutes at room temperature in the dark. The sample was rinsed again with 4X SSC before an anti-photobleaching solution consisting of 50 mM Tris HCl (Invitrogen #15568-025), 1 mM Trolox (Sigma-Aldrich #238813), 10% glucose (Sigma-Aldrich #G7528-250G), 1X catalase (Sigma-Aldrich #C3155-50MG), 1 mg/mL glucose oxidase (Sigma-Aldrich #G2133-10KU), and 4X SSC was applied for imaging. RNA integrity was confirmed by confocal microscopy with signal colocalization in the 647 nm and 561 nm fluorescent channels. After RNA integrity was confirmed, automated RNA seqFISH+ imaging was performed on the Spatial Genomics GenePS following the manufacturer’s instructions. For each slide, a maximum of 100 mm^2^ area was imaged with 3 Z-stacks with 1.5 μm spacing. Exposures were set to 350 ms for the 647nm, 561 nm, and 488 nm channels and 150 ms for the 405 nm channel.

### DNA seqFISH+ Hybridization and Imaging and Immunofluorescence

After RNA seqFISH+ imaging was complete, the sample was stored at 4°C in 4X SSC before processing for DNA seqFISH+. First, the sample was washed 3 times with 1X PBS. Next, RNase A/T1 mix (Thermo Scientific #EN0551) was diluted 1:100 in 1X PBS and incubated on the sample for 1 hour at 37°C. After RNase incubation, the sample was washed 3 times with denaturation buffer consisting of 50% (v/v) formamide in 2X SSC. Next, the flow cell was sealed, and the sample was incubated in denaturation buffer on a 90°C heat block for 7 minutes. DNA seqFISH+ probe hybridization solution was prepared to contain 0.75 nM/oligo barcoded DNA seqFISH+ probes, 1 nM/oligo additional gene DNA probes, 10 nM/oligo *EGFR* probes, 1 nM/oligo repetitive element probes, 37.5% (v/v) formamide, and 4X SSC. The DNA probe hybridization solution was warmed at 37°C then heated for 3 minutes at 90°C. After the sample denaturation was complete, the sample was washed 3 times with 4X SSC then the DNA probe hybridization solution was added at 90°C. Next, the flow cell was sealed and the sample was incubated at 37°C for 4–7 days in a humidified chamber. After incubation, the sample was washed 3 times for 30 minutes with 42.5% (v/v) formamide, 0.1% triton X-100, 4X SSC wash buffer at 37°C. Afterwards, the sample was washed 3 times with 0.1% triton X-100 in 1X PBS (PBST). Next, the hybridized probes were circularized by ligation of a splint oligonucleotide complementary to the 5’ end of forward primer and the 3’ universal sequence. First, the sample was washed with 1X Quick Ligase Buffer (New England Biolabs #B2200S). Next, the sample was incubated at room temperature for 1 hour with 100 μM splint oligonucleotide (Integrated DNA Technologies) and 1:10 Quick Ligase (New England Biolabs #M2200S) in 1X Quick Ligase Buffer. Afterwards, samples were washed 3 times with PBST and 3 times with 4X SSC.

DNA seqFISH+ imaging was performed on the Spatial Genomics GenePS system. For each imaging round, 220 μL readout solution was prepared using 50 nM per fluorescent readout in 10% (w/v) ethylene carbonate (Sigma-Aldrich #E26258), 100 mg/mL 6,500–10,000 MW dextran sulfate (Sigma-Aldrich #D4911), and 4X SSC. Automated DNA seqFISH+ imaging was performed on the Spatial Genomics GenePS following the manufacturer’s instructions. For each slide, a maximum of 100 mm^2^ area was imaged with 23 Z-stacks with a 0.3 μm spacing or 10 Z-stacks with 0.7 μm spacing. Exposures were set to 350 ms for the 647nm, 561 nm, and 488 nm channels and 150 ms for the 405 nm channel.

### Image Processing

Images were processed using the SG Navigator software (Spatial Genomics, Inc.). Images were aligned and background subtracted. Cell boundaries were identified using nuclear DAPI signal with nuclear mask dilation. Dot detection was performed with a combination of manual and automatic thresholding per the companies’ recommendations. Finally, dots were decoded according to the barcoding scheme for RNA and DNA seqFISH with a false discovery rate (FDR) of 5%.

### ecDNA Analysis

ecDNA was quantified on a single cell basis and defined as a gene locus detected further than 1 μm from its typical chromosome territory (*PDGFRA*, chr4; *EGFR*, chr7). The number of ecDNA loci was counted per nucleus and compared between samples using the Mann-Whitney U test.

### Pairwise Spatial Distance Analysis for DNA Loci

Distance calculations were performed on a single cell basis. For each DNA locus, the minimum Euclidean distance to all other DNA loci was calculated. The minimum distance was converted to contact probability using the inverse power law. Next, the contact probability maps were corrected using a method adapted from a Hi-C data correction method described by Imakaev et al.^66^. First, the per-locus row and column biases in normal and tumor cell types were calculated by averaging across all cells of that cell type. The shared component of these biases was estimated via linear regression of the tumor per-locus averages onto the normal per-locus averages. A geometric mean of the predicted shared biases from each matrix was then used as the correction factor. Each matrix element was divided by the outer product of the shared row and column bias vectors, effectively removing the correlated stripe patterns. The corrected matrices were then rescaled to preserve the original median contact frequency. A second round of correction was then performed to remove the effects of copy number alterations in tumor cells and highlight interchromosomal interactions. The tumor cell distance matrix was divided by its per-locus row and column biases then rescaled to the average bias profile of the normal cell matrix.

### Single Nucleus RNA Sequencing

Single nucleus RNA sequencing was performed on 10 samples for which both RNA and DNA seqFISH+ imaging was performed. Fresh frozen glioma tissue was equilibrated in a cryostat at −20°C for 30 minutes prior to sectioning. Tissue was affixed to a cryostat specimen chuck using OCT embedding medium and sectioned at 60 μm. A minimum of 15 mg of tissue per sample was sectioned, collected in a microcentrifuge tube, and stored at −150°C prior to processing.

To prepare single nuclei for sequencing, samples were first lysed in 1 mL nuclear extraction buffer (NEB) comprised of nuclei extraction buffer (Miltenyi Biotec #130128024) containing 0.2 U/μL RNase inhibitor (Millipore Sigma #3335402001). Lysate was mixed with an additional 1 mL NEB then passed through a 200 μm cell strainer (pluriSelect #431020040). The strainer was washed with an additional 2 mL NEB, and the flowthrough was centrifuged at 300 x g for 5 minutes at 4°C. The supernatant was aspirated, and the pellet was resuspended in 1.5 mL nuclear separation buffer (NSB) comprised of 4% BSA (Miltenyi Biotec #130091376), 0.2 U/μL RNase inhibitor, 14% (v/v) nuclei extraction buffer, and 1X PBS. The resuspended nuclei were then passed through a 40 μm cell strainer (pluriSelect #431004060), and the flowthrough was centrifuged at 300 x g for 5 minutes at 4°C. After aspirating the supernatant, the pellet was resuspended in 900 μL NSB and mixed with 100 μL anti-nucleus MicroBeads (Miltenyi Biotec #130132997). The nuclei were incubated with the MicroBeads for 15 minutes at 4°C in a tube rotator set to 4 rpm. After incubation, the labeled nuclei were mixed with 2 mL NSB and magnetic separation was performed on a LS column (Miltenyi Biotec #130042401) using a QuadroMACS Separator (Miltenyi Biotec #130090976) following the manufacturer’s protocol. After performing magnetic separation, the nuclei-enriched solution was centrifuged at 300 x g for 5 minutes at 4°C, and the pellet was resuspended in 100 μL calcium and magnesium-free PBS (Gibco #10010023) containing 1% BSA, 0.2 U/μL RNase inhibitor, and 1 mM DTT (Sigma-Aldrich #646563). The concentration and purity of the nuclear preparation was evaluated by mixing 20 μL resuspended nuclei with 1 μL 10 μg/mL Hoechst 33342 (Biotium #40046) and counting on a hemocytometer.

Single nucleus RNA sequencing libraries were prepared using the Chromium GEM-X Single Cell 3’ Reagent Kit v4 (10X Genomics #1000691) following the manufacturer’s protocol. The completed libraries were quantified using Qubit dsDNA HS (High Sensitivity) Assay Kit (Invitrogen #Q32851), and fragment size distribution was assessed by Bioanalyzer High Sensitivity DNA Kit (Agilent #50674626). Libraries were sequenced on a NovaSeq X Plus at the University of California San Francisco Center for Advanced Technology (UCSF CAT). Sequencing files were processed using Cell Ranger^66,67^ v10.0.0 (10X Genomics) prior to downstream analysis.

### Inferred CNA Analysis

Inferred CNA analysis from single nucleus RNA sequencing data was performed using a custom Python implementation of the inferCNA approach^30^. Using cell type annotation from RNA clustering performed using Seurat^68^, single cells were classified as tumor cells and normal cells, with the mean of the normal cells used as the baseline for CNA inference. First, the data was filtered to remove cells with fewer than 200 genes detected, greater than 8,000 genes detected or greater than 20% mitochondrial read fraction. Genes were restricted to autosomes (chr1–chr22). Genes found in fewer than 3 cells were also removed. Filtered counts were normalized to 10,000 counts per cell and log1p-transformed prior to CNA inference. Log-normalized expression values were mean-centered per gene and per cell, and value were clipped to +/− 3 to limit the influence of outlier expression values. The resulting matrix was smoothed along the linear genomic gene order using a pyramidal moving average window of 101 genes. The mean smoothed profile of the normal cells was subtracted, and a per-cell CNA score was defined as the standard deviation of the smoothed, baseline-relative expression profile across all genes.

To compare tumor cell autosome CNAs detected by DNA seqFISH and inferred CNAs from snRNAseq, each 3 Mb DNA seqFISH genomic region was compared to the average inferred CNA of genes within each region. CNAs were averaged across all tumor cells from each method and a Pearson correlation coefficient was calculated.

### Spatial transcriptomic data processing and cohort definition

Counts matrix was normalized to 10,000 transcripts per cell and then log1p-transformed. After quality filtering the dataset comprised 695,099 cells by 1,150 genes across 40 tissue sections, with cell centroids in micrometres.

Analyses of malignant transformation used a discovery cohort of 16 sections from 13 patients (190,951 cells): 10 sections from low grade glioma patients who subsequently progressed to high grade (future-MT: 2122, 2411_1, 2411_2, 2683, 2789, 2830, 3209, 3557, 4494_1, 4494_2) and 6 from patients who remained stable (future-stable: 3364, 4523, 4668, 4970, 5272_1, 5272_2). Sections 2411_1/2411_2, 4494_1/4494_2 and 5272_1/5272_2 are pairs from the same patient. All cross validation folds and all patient level statistics treat each patient as a single unit.

Cell types were called from the RNA data as described above (see “RNA Cell Type Clustering”). For all cell pair analyses the three tumor populations (OPC-like, OPC-like proliferating and AC-like) were merged into a single tumor label unless a subtype was explicitly required, giving ten grouped cell types.

### Spatial neighbourhood and cell-pair conventions

Unless stated otherwise, two cells were considered spatial neighbors when the Euclidean distance between their centroids was ≤ 50 micrometers. Neighbourhood graphs were constructed independently within each tissue section, so no edge crosses a section or a patient. A gene was expressed in a cell when at least one raw transcript was assigned to it, this threshold was applied to raw counts and never to normalized values. For an ordered cell type pair (sender to receiver), a cell pair is an ordered pair of one sender cell and one receiver cell within 50 micrometers of each other, so that a sender with k receiver type neighbors contributes k cell pairs. Where sender and receiver cell types are identical, the focal cell was excluded from its own neighbour set in the differential expression analysis and retained in the risk score analysis.

### Ligand receptor co-expression on spatial cell pairs

We define cross-expression of a ligand–receptor (LR) pair on a cell pair as the joint event that a sender cell expresses the ligand and a spatially adjacent receiver cell expresses the receptor. For an ordered cell type pair (A to B) and a gene pair (ligand gene 1 in A, receptor gene 2 in B) we computed, within each section, the binary 50 micrometer adjacency matrix S between all A cells and all B cells, binarized expression at ≥ 1 raw transcript to give indicator vectors a and b, and defined the cell pair co-expressed fraction as the number of within 50 micrometer A and B cell pairs that are double positive for (gene 1, gene 2), divided by the total number of within 50 micrometer A and B cell pairs. This gives one value per (cell type pair, gene pair, section), and is the quantity plotted in the box plots of Fig. 4B and 4E. Where sender and receiver cell types are identical the adjacency matrix includes the diagonal, so a cell is its own neighbour. It contributes one cell pair to the denominator and, if it expresses both genes, one to the numerator as well. The two marginals in the accompanying scatter plots are computed at the cell rather than the cell pair level: the fraction of sender cells expressing the ligand among sender cells that have at least one receiver type neighbor within 50 micrometer, and the analogous fraction for receivers.

The statistic was computed over 154 ordered cell type pairs: all 100 ordered pairs of the ten grouped cell types, plus the 27 pairs from each non-tumor type to each of the three tumor subtypes and the 27 reciprocal pairs, so that subtype resolved interactions could be examined where needed. A cell type pair was tested only if at least three future MT and three future stable sections contained cell pairs of that type. Cross-expression fractions were compared between future-MT and future-stable sections with a two-sided Mann–Whitney U test using the exact null distribution where sample size permits; with 10 versus 6 sections the smallest attainable two-sided P value is 2.5 × 10⁻⁴. The same metric was additionally computed across all 40 sections on the 1,150-gene dataset.

### Cell pair discovery and permutation false discovery rate

Candidate gene pairs were restricted to the NicheNet ligand–receptor network intersected with the 1,150-gene panel, giving 769 pairs, to which ENPP2 to LPAR1 was added explicitly because it is absent from that database, for a total of 770 ligand receptor pairs spanning 257 ligands and 250 receptors.

The test was always defined between two arms - future MT and future stable. A (cell type pair, gene pair) hypothesis was eligible for testing when at least three future MT and at least three future stable sections had a non zero co-expressed fraction and the larger of the two arm means exceeded 0.05; applying this gate left 19,682 testable hypotheses. Each was tested as described above, and hypotheses reaching two-sided Mann–Whitney P < 10⁻³ were called significant.

The false discovery rate was estimated by permutation, because the hypotheses are strongly dependent, the same gene pair is tested across many cell-type pairs and the same cells contribute to many hypotheses. Holding the eligibility gate fixed at the set defined by the real labels, we randomly permuted the future MT/future stable labels across the 16 sections 100 times and recomputed every Mann–Whitney test on that fixed hypothesis set. The FDR at threshold alpha was estimated as the mean number of permutation rejections divided by the number of observed rejections. At alpha = 10⁻³, *R*_obs_ = 42 and *E*[*V*_null_] = 6.5, giving FDR = 15.4%.

Of the 42 observations, 18 resolved a single tumor subtype rather than the merged tumor population and were removed so that every reported interaction refers to the same tumor definition, leaving the 24 cell pairs used throughout; subsetting does not lower the false discovery rate, which the 24 inherit from the full observation set. These 24 comprise 12 distinct ligand receptor axes, of which ENPP2 to LPAR1 accounts for nine across different cell type combinations, and include two reciprocal duplicates (CLDN11 with CLDN11 and PECAM1 with PECAM1), giving 22 distinct cell populations.

### Validation cohort and sequencing depth equalization

An independent validation cohort of four patients (one future-MT, three future-stable) was profiled with a reduced 130-gene seqFISH panel containing cell type markers together with ENPP2 and LPAR1, yielding 19 regions of interest. Cell types were assigned by reference mapping onto the cell type definitions of the discovery cohort so that both cohorts use the same labels.

Co-expressed fractions were computed exactly as in the discovery cohort (50 micrometer, ≥ 1 raw transcript) and compared between future MT and future stable at the level of individual regions of interest with a two sided Mann–Whitney U test. 13 of the 24 discovered cell pairs are testable on the 130-gene panel.

Because the two cohorts were imaged with different panels and differ systematically in transcripts detected per cell, and because the co-expressed statistic depends on detection efficiency for both partners, the validation was performed on raw counts and, in parallel, after equalizing RNA seqFISH count depth. For depth equalization, transcripts were pooled within each region of interest and resampled without replacement to the smallest per-cell median depth across regions, so that every region contributes the same number of transcripts per cell. Both count modes are reported.

### Signaling specific differential expression

To ask which genes a cell changes when it participates in a signaling cell pair, each host cell type was split into “cross-expressing” and “rest” populations with respect to a given partner cell type. A host cell was called “cross-expressing” when it expressed its own gene of at least one of the 24 discovered ligand receptor pairs connecting the two cell types, in either direction, and had at least one partner type neighbor within 50 micrometer expressing the corresponding partner gene, with the focal cell excluded from its own neighbor set. Because “cross-expressing” cells are by construction a subset of the cells merely co-localized with the partner, three complementary rest definitions were evaluated: all other host cells; host cells co-localized with the partner but not “cross-expressing”, which holds proximity fixed and isolates the co-expression effect; and host cells with no partner-type neighbor within 50 micrometer. The published contrast uses the first of these — rest = all other host cells of that type.

Differential expression was computed as a two-stage per section statistic rather than by pooling cells, so that sections rather than cells are the unit of replication. Within each section the mean expression difference between “cross-expressing” and “rest” cells was computed for every gene, requiring at least 10 cells in both groups for that section to contribute; the arm-level effect is the equal weight mean of the usable per section differences. The number of usable sections therefore varies by panel and by arm, for microglia contacting tumor cells, for instance, all 10 future MT but only 5 of the 6 future stable sections contribute. Significance was assessed in two ways, both computed and reported: a two-sided Mann–Whitney U test of interacting versus rest cells within each section, combined across sections. P values were adjusted across genes by the Benjamini– Hochberg procedure.

Genes were restricted to those specific to the host cell type, defined as the host mean expression divided by the sum of the host mean and the largest other-cell-type mean, thresholded at > 0.5. The analysis was run on raw counts, with the host-specificity gate computed on the same matrix. The diagonal representation of Fig. 4H plots the future stable effect against the future MT effect for each gene, so that genes lying on the diagonal have a contact response of equal magnitude in the two arms.

### Cell Interaction Foundation Model and microenvironment graph

The Cell Interaction Foundation Model (CIFM) is a graph neural network that predicts a cell’s transcriptional state from the states and spatial coordinates of the cells around it. We used the 100M-parameter checkpoint, pre-trained on 23 million microenvironments from healthy and diseased human tissues.

Each tissue section is represented as a graph over cell centroids. The neighborhood radius is derived from the data rather than fixed. A radius graph with self-loops was then built at that radius within each section, and the microenvironment of a cell was defined as its 2-hop subgraph. During both training and inference the central cell’s own expression is masked from the model input, so a cell’s state is predicted strictly from its neighbors.

The pre-trained model was fine-tuned on our complete glioma seqFISH+ dataset (695,099 cells across all 40 sections, with section identity as the batch key) for 20 epochs at a learning rate of 1 × 10⁻⁶ and batch size 2, using the model’s self-supervised expression-reconstruction objective. The gene panel was mapped onto the model’s gene vocabulary by Ensemble identifier and unmatched channels were dropped. Fine-tuning uses no clinical labels.

### Forward tissue simulation

CIFM was used as a generative forward simulator by iterating its update rule. At each step every cell in a section was re-predicted from its 2-hop microenvironment and the predicted expression vector replaced that cell’s state for the next step; cell coordinates and graph topology were held fixed throughout, so the simulation changes transcriptional state alone and never cell position, cell number or tissue architecture. Each section was simulated independently for 30 steps, with step 0 defined as the measured data, producing a 1,150-dimensional expression trajectory for every cell. No gene was perturbed, so the trajectory is the model’s unforced baseline.

### Progression risk classifier

A random-forest classifier (300 trees, maximum depth 10, scikit-learn, seed 42) was trained to predict the clinical label of a cell’s section from its 1,150-dimensional expression vector. Cross-validation folds were assigned by patient across the 13 discovery patients, so all sections from a patient were held out together and no cell from a test patient contributed to training. Within each fold the training pool was drawn balanced across training patients, an equal number of cells per patient, subsampled to the smallest patient, and was assembled from all 31 simulation steps so that a single classifier applies at every step; this gives 1.5–2.4 × 10⁶ training rows per fold, of which the measured data (step 0) contributes 3.2%.

Every cell was scored at every step by the fold that had not seen its patient; the risk score is that out-of-fold predicted probability of the future-MT class. Per-section risk was summarized as the mean over cells and compared between arms with a two-sided exact Mann–Whitney U test, at step 0, at the step of maximal separation and at step 30. Spatial risk maps (Fig. 6C) plot each cell at its measured coordinates colored by its risk score on a fixed [0, 1] scale, so that panels are comparable across steps and sections.

### Association of the risk score with signaling cell pairs and gene expression

To ask whether cells engaged in signaling cell pairs carry higher risk, tumor cells were split into “cross-expressing” and “rest” populations. A tumor cell was called “cross-expressing” when it expressed its own gene of the pair and had at least one partner-type neighbor within 50 micrometer expressing the partner gene; all other tumor cells formed the comparison group. Risk scores on the measured (step 0) data were compared between the two groups with a two-sided Mann–Whitney U test and Cliff’s delta, where positive values indicate that contacting cells carry higher risk. Because pooling cells across sections confounds the comparison with section identity, the same statistic was recomputed within each section and summarized by its median with a sign test across sections; both are reported. The comparison was run pooled across the cohort and separately within the future-MT and future-stable arms.

To relate gene expression to risk without a between-section comparison, Spearman correlations between a gene’s expression and the per-cell risk score were computed within each section among cells of a single type and then averaged across sections, and tested against zero by a two-sided Wilcoxon signed-rank test on the per-section correlations.

### Sensitivity and coverage down-sampling

To ask how much detection sensitivity and how much imaged area the cell pair discovery requires, the measured data were downsampled along two independent axes and the cell-pair statistic recomputed.

For the transcript count axis, all transcripts of a section were pooled and a fraction 1/f drawn without replacement by a multivariate hypergeometric sample, for f in {2, 3, 5, 10, 30, 50}, corresponding to 50%, 33%, 20%, 10%, 3% and 2% of the original counts; this preserves each section’s relative gene composition while lowering per cell detection, simulating a lower sensitivity platform. For the imaged-area axis, each section was partitioned into square tiles of 256 units and a random subset retained at coverages of 80%, 60%, 40% and 20%; cells outside the retained tiles were removed before the neighborhood graph was rebuilt, so that cells and their spatial context are lost together as they would be with a smaller imaged region. One hundred independent draws were generated at every level.

For each draw the 50 micrometer cell-pair cross-expressed fraction was recomputed and re-tested between future MT and future stable sections with the two-sided Mann–Whitney U test. At every draw the entire ligand receptor space 48,910 cell-type-pair times gene-pair entries over the 84 cell-type pairs with sufficient sampling in both arms, was re-gated and re-tested under the full discovery criterion, so that a cell pair may enter the significant set as well as leave it. The plotted quantity is therefore a systematic evaluation of everything significant at that level of downsampling rather than the survival of a fixed list, and it equals 24 at full data because that is the criterion which produced the 24 pairs.

## ACKNOWLEDGEMENTS

We thank Maydelle Lorenzo for assistance with automated seqFISH imaging and technical support, Chee-Huat Linus Eng and Kirsten Frieda for technical support, Yujing Yang for assistance with DNA seqFISH design and analysis, Yodai Takei for antibody conjugation and assistance with DNA seqFISH design and analysis, Jonathan White for consulting on analysis, Diane Trout for IT support, Shivani Baisiwala for electronic health record review, and Lu Sun for assistance with inferred CNA analysis.

